# Multimodal hierarchical classification of CITE-seq data delineates immune cell states across lineages and tissues

**DOI:** 10.1101/2023.07.06.547944

**Authors:** Daniel P. Caron, William L. Specht, David Chen, Steven B. Wells, Peter A. Szabo, Isaac J. Jensen, Donna L. Farber, Peter A. Sims

## Abstract

Single-cell RNA sequencing (scRNA-seq) is invaluable for profiling cellular heterogeneity and dissecting transcriptional states, but transcriptomic profiles do not always delineate subsets defined by surface proteins, as in cells of the immune system. Cellular Indexing of Transcriptomes and Epitopes (CITE-seq) enables simultaneous profiling of single-cell transcriptomes and surface proteomes; however, accurate cell type annotation requires a classifier that integrates multimodal data. Here, we describe <u>M</u>ulti<u>Mo</u>dal <u>C</u>lassifier <u>Hi</u>erarchy (MMoCHi), a marker-based approach for classification, reconciling gene and protein expression without reliance on reference atlases. We benchmark MMoCHi using sorted T lymphocyte subsets and annotate a cross-tissue human immune cell dataset. MMoCHi outperforms leading transcriptome-based classifiers and multimodal unsupervised clustering in its ability to identify immune cell subsets that are not readily resolved and to reveal novel subset markers. MMoCHi is designed for adaptability and can integrate annotation of cell types and developmental states across diverse lineages, samples, or modalities.

## INTRODUCTION

Recent advances in high-dimensional profiling of single cells, most notably by single-cell RNA sequencing (scRNA-seq), have transformed our ability to define the function and heterogeneity of cell populations in diverse biological systems^1–4^. Because many of the features that define cell types are not captured by scRNA-seq, multimodal single-cell technologies have been developed, including Cellular Indexing of Transcriptome and Epitopes by sequencing (CITE-seq)^5^, RNA Expression And Protein sequencing assay (REAP-seq)^6^, and Antibody sequencing (Ab-seq)^7^ for simultaneous profiling of surface proteomes and transcriptomes. Integrating these high-dimensional modalities to identify cell subsets, developmental states, and other cellular properties with high fidelity across disparate datasets remains a challenge.

Downstream of raw data processing, the first and most important basic analytical step for single-cell sequencing is classification of individual cells, often in terms of canonical subsets. The vast majority of analytical tools, from differential expression analysis to trajectory inference to statistical modeling with clinical co-variates, depend on this crucial initial step. Many analytical tools have been developed for the cellular annotation of scRNA-seq data (reviewed in Ref.^8^ and Ref.^9^). Unsupervised clustering is commonly used to divide cells into groups with similar expression profiles^10,11^. This approach has proven invaluable for characterizing cellular heterogeneity^12^, and has been adapted for CITE-seq datasets^13,14^. However, the number, type, and identity of clusters can be difficult to compare across studies^12,15^.

On the other hand, supervised machine learning methods can address these drawbacks by incorporating reference data or marker-based definitions of cell types^16^. Examples of such tools include CellTypist^17^, ImmClassifier^18^, HieRFIT^19^, and Garnett^20^. CellTypist uses logistic regression and stochastic gradient descent for automated annotation based on a reference atlas^17^. To overcome the many, sometimes overlapping, annotations across atlases, it includes an option called “majority voting” where users can elect to perform conventional clustering and annotate all cells in each cluster with the most dominant classification. ImmClassifier addresses the reverse problem—existing reference datasets for cell types of interest (i.e. for immune cells) may not be pre-annotated with the desired granularity or may not include all the necessary cell subsets^18^. Thus, it first annotates input data with multiple random forest classifiers trained on independent reference datasets and then generates a consensus classification using a deep neural network. To improve annotation accuracy of highly similar subsets, HieRFIT utilizes a hierarchical design, implementing multiple random forest classifiers to progressively annotate cell types^19^. Together these reference-based methods enable cross-study comparisons^16,21^, but require reference atlases which are not available for all tissues and contexts^22–24^. Reference-free annotation methods address these shortcomings, instead relying on knowledge-based marker definitions^20,25^. Garnett, for example, classifies scRNA-seq data using a hierarchy of cell types defined by user-supplied markers and a regularized elastic net generalized linear model for training^20^. Overall, existing tools have successfully leveraged a variety of machine learning algorithms, both reference-based and marker-based cell type definitions, and hierarchical designs for cell type classification, all with exclusively RNA-level features.

Here, we developed a supervised approach for cell type annotation of CITE-seq data, designated <u>M</u>ulti-<u>Mo</u>dal <u>C</u>lassifier <u>Hi</u>erarchy (MMoCHi) that incorporates surface protein and transcript features for marker-based, reference-free classification. The development of reference-free, multimodal classifiers to analyze single-cell data is of particular importance for studying the immune system. Immune cells comprise multiple disparate lineages—each of which can be subdivided into functionally distinct, but closely related subsets that represent various developmental and activation states^26^. Decades of research have delineated numerous immune cell subsets across the lymphocyte and myeloid lineages, often defined by surface phenotypes that shape our biological questions^26–29^ and are challenging to annotate by transcriptome alone^18,30–33^. Thus, to benchmark this tool against other annotation methods, we sorted and profiled T cell subsets using CITE-seq and demonstrate improved performance by MMoCHi over the alternative methods described above, particularly in the annotation of subsets with highly similar expression profiles. We applied MMoCHi to produce integrated annotations for a cross-tissue CITE-seq atlas of diverse immune cell populations isolated from complex samples. Extracting important features from MMoCHi’s underlying random forests revealed highly interpretable learned representations of cell types, which we used to identify novel markers for distinguishing transcriptionally similar T cell subsets. Finally, we demonstrated the versatility of MMoCHi by extending to applications beyond CITE-seq. This includes identification of transformed tumor cells from joint profiling of aneuploidy and gene expression and annotation of *in situ* spatial profiling (10x Genomics Xenium) by joint classification with gene expression and morphological features. Together, MMoCHi readily enables multimodal cell type annotation based on marker genes and proteins, is designed for applicability to CITE-seq of any cell lineage or sample type and shows promise for extension to forthcoming modalities such as *in situ* spatial profiling of complex tissues.

## RESULTS

### Algorithm Overview

The MMoCHi algorithm uses a hierarchy of random forest classifiers trained on gene expression (GEX) and antibody-derived tags (ADTs) for cell type classification of CITE-seq data (Fig. 1). Prior to classification, ADT expression is batch-corrected using landmark registration, as previously applied to flow cytometry^34^ and CITE-seq^35^ (see Methods). We identify populations exhibiting negative (background) and positive ADT expression and apply warping functions to align their midpoints (landmarks) across batches, to effectively integrate CITE-seq expression (Fig. 1a). MMoCHi then classifies cell types based on a user-supplied hierarchy of cell subsets paired with marker-based definitions (Fig. 1b). At each classification node, high-confidence members of each subset are identified using manual thresholds on user-provided gene and protein markers. A random forest is then trained on a representative set of these high-confidence cells and used to annotate all cells—including those not labeled by high-confidence thresholding (Fig. 1b; Extended Data Fig. 1). Once trained, classifiers can be interrogated for features important for cell classification or applied to extend cell type annotation to other datasets.

**Figure 1.**
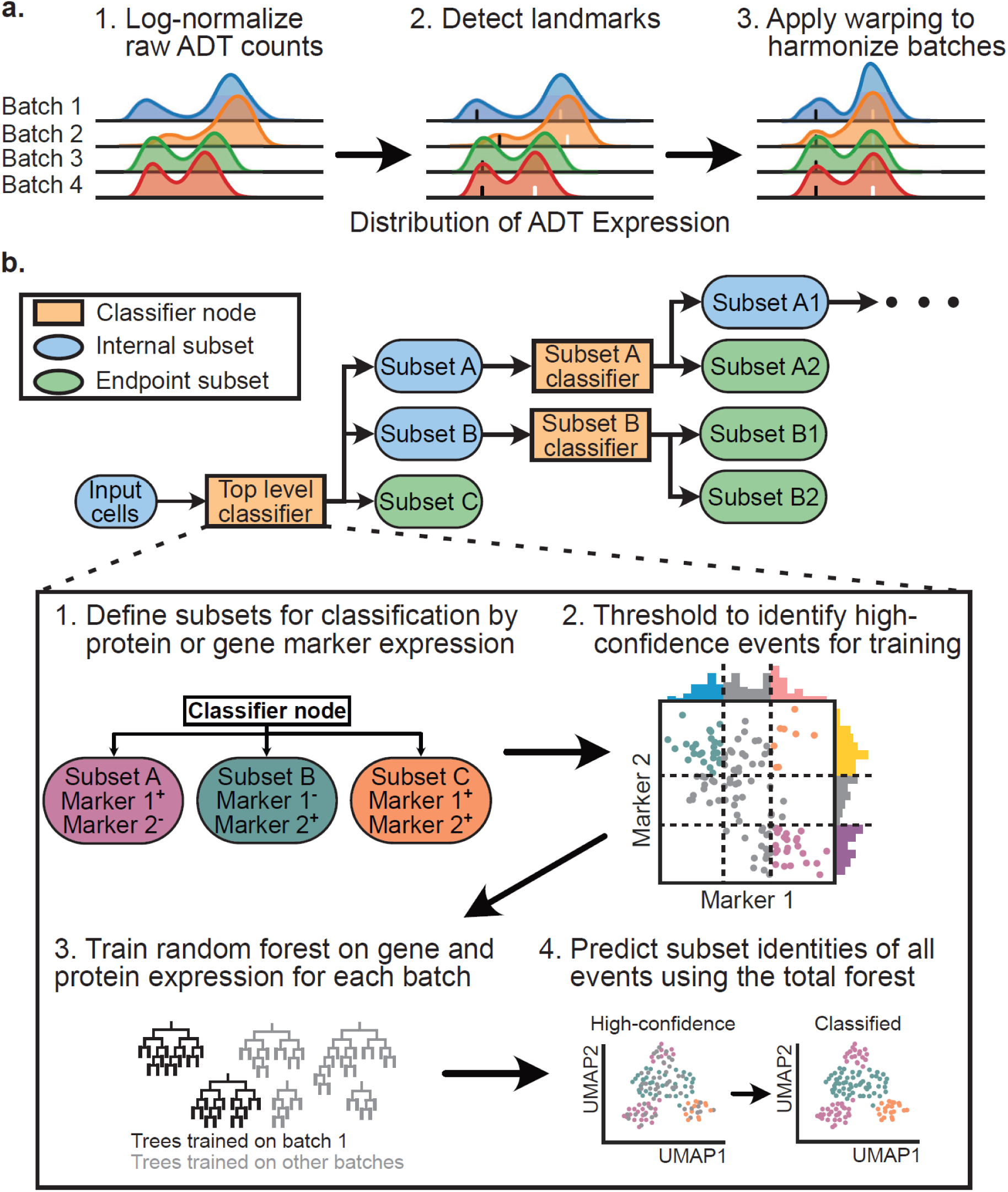
Schematic showing general workflow of MMoCHi. a. Batch correction of Antibody Derived Tag (ADT) expression was performed by landmark registration on log-normalized counts (1). After detection of positive (white ticks) and negative (black ticks) peaks in each batch (2), a warping function was applied to align these landmarks across batches (3). b. MMoCHi hierarchy demonstrating the classification workflow. User supplies a hierarchy of cell subsets, and marker definitions for each subset (1). Thresholding is performed to select high-confidence events for each subset (2). A portion of these high-confidence events are used to train a random forest (3; see Extended Data Fig. 1). Finally, the trained random forest is used to predict subset identities of all events from the parent subset (4). This process is repeated for each classifier node within the hierarchy until all cells are classified to endpoint subsets.

### Superior annotation of closely related immune cell subsets by MMoCHi

MMoCHi is designed to improve the annotation of subsets with highly similar transcriptomic profiles compared to conventional manual annotation of clusters or methods that rely exclusively on mRNA profiles. We chose to test its performance using T cell subsets, which are well-defined by surface marker expression and functional readouts^27,28^, but challenging to annotate by scRNA-seq alone. For example, CD4^+^ and CD8^+^ T cells are often imperfectly resolved, partially due to low *CD4* transcript expression^18,19,31^. Moreover, conventional human T cells are delineated into subsets based on differentiation state and migration capacity into naive T cells (CCR7^+^ CD45RA^+^ CD45RO^-^) and memory T cells, which comprise central memory (T_CM_; CCR7^+^ CD45RA^-^ CD45RO^+^), effector memory (T_EM_; CCR7^-^ CD45RA^-^ CD45RO^+^), and terminally differentiated (T_EMRA_; CCR7^-^ CD45RA^+^ CD45RO^-^) subsets^27,28^. Distinguishing naive T cells from T_CM_ and T_EM_ from T_EMRA_ by transcriptome alone is challenging^1,18,30,31^. We sorted and performed CITE-seq on seven T cell subsets (CD4^+^ naive, CD4^+^ T_CM_, CD4^+^ T_EM_, CD8^+^ naive, CD8^+^ T_CM_, CD8^+^ T_EM_, and CD8^+^ T_EMRA_) and monocytes (Fig. 2a; Extended Data Fig. 2a; Supplementary Tables 1, 2). CCR7 staining by CITE-seq is suboptimal^5^, so we used CD62L which has high concordance in human blood^27,28^. To eliminate batch effects, we performed CITE-seq antibody staining before sorting and labeled sorted populations with hashtag antibodies prior to pooling all samples for library preparation and sequencing. Hashtagged populations then served as known references to evaluate classification.

**Figure 2.**
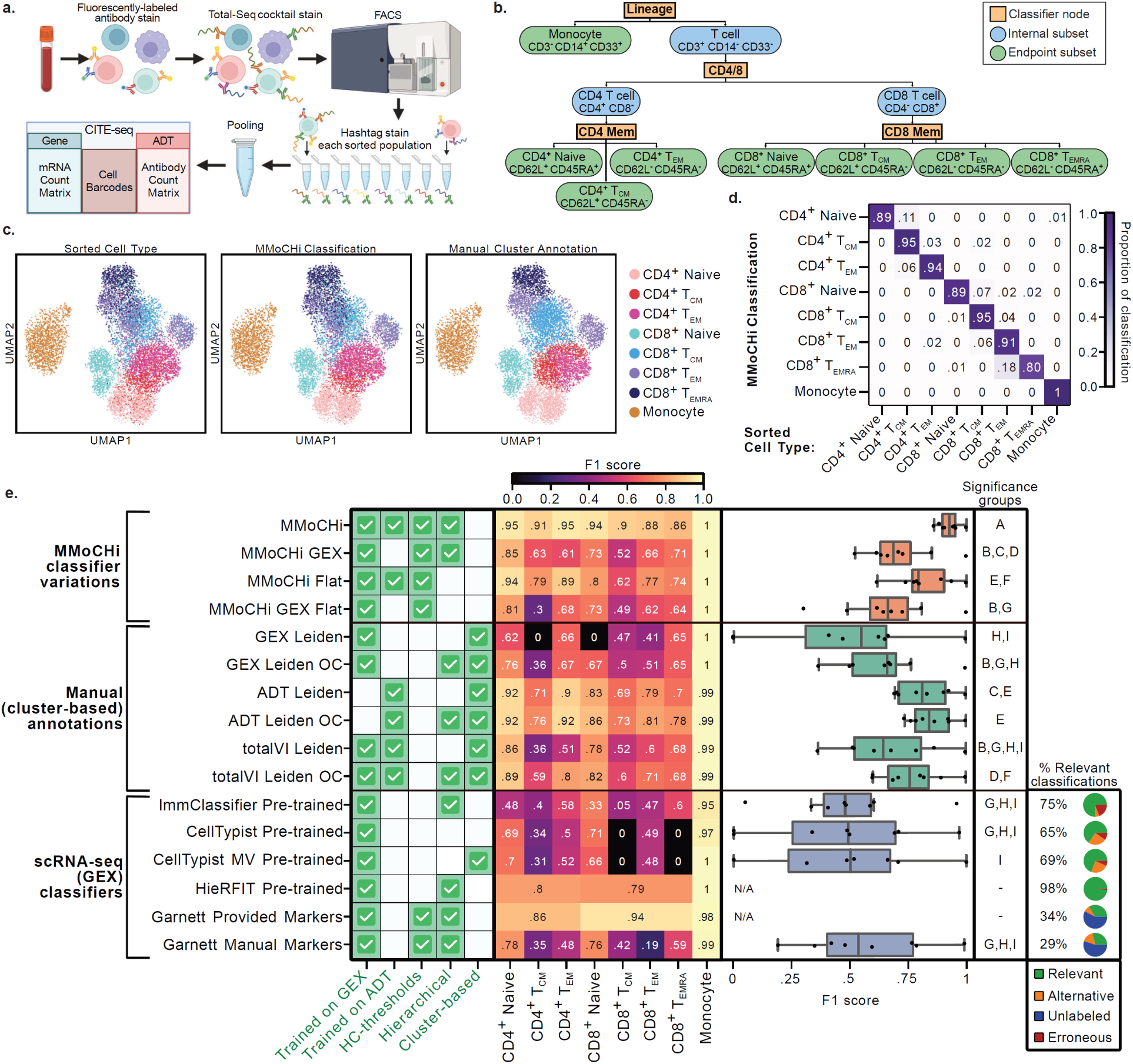
MMoCHi classification of predefined T cell subsets outperforms other annotation methods. a. PBMCs were stained with fluorescently-labeled antibodies and a cocktail of oligo-tagged antibodies for sequencing. Seven T cell subsets and monocytes were sorted, hashtagged, pooled, and sequenced. b. MMoCHi hierarchy defining subsets using the same markers used for sorting. c. UMAPs of totalVI latent space colored by sorted cell type (identified by hashtag oligo; HTO), MMoCHi classification, or manual cluster annotations. d. Row-normalized heatmap comparing MMoCHi classification to sorted cell type. Color represents proportion of cells in each MMoCHi classification from each sorted subset. e. Performance comparison using F1 scores, calculated for each cell subset using HTO-derived sorted cell type labels as truth. F1 scores for each method were aggregated in box and whisker plots (right). Features of each method are labeled for context (left) using green checks: “Trained on GEX/ADT”—whether gene expression (GEX) or antibody derived tag (ADT) expression data were used during model training. “HC-thresholds”—whether high-confidence thresholds were used for training data selection. “Hierarchical”—whether annotation was performed on multiple levels. “Cluster-based”—whether annotations were applied to unsupervised clusters. For scRNA-seq classifiers, the percent of classifications that were relevant to this analysis, were alternative (but potentially valid) classifications, were unlabeled, or were entirely erroneous are shown using pie charts (far right). Only relevant classifications were included for calculation of F1 scores. Statistical significance was calculated using a Friedman rank-sum test, matched by subset, followed by multiple comparisons testing using paired Wilcoxon signed-rank tests and FDR correction. Significance groups for each method are shown by lettering, where methods not sharing a letter are significantly different (p < 0.05). Methods that could not label T cell memory subsets were excluded from significance testing and the merged efficacy of CD4^+^ or CD8^+^ T cell classifications were displayed instead. MV, Majority voting; T_CM_, central memory T cell; T_EM_, effector memory T cell; T_EMRA_, terminally differentiated effector memory T cell. Schematic in (a) created with BioRender.com

All sorts were of high purity (>91%; Supplementary Table 3), and subsets reflected expected ADT marker expression (Extended Data Fig. 2b). We devised and applied a MMoCHi hierarchy with the same protein markers used for sorting (Fig. 2b, c; Supplementary Table 4). For comparison to unsupervised approaches, we used totalVI^14^ to calculate a multimodal latent space, then performed clustering and manual annotation (Fig. 2c). The high concordance between sorted cell type and MMoCHi classification compared to manual annotation is visually evident from UMAPs in Fig. 2c. Indeed, MMoCHi classification had 92% agreement with sorted labels, while manual annotation had only 70% agreement (Fig. 2c,d; Supplementary Table 5). MMoCHi classified populations showed expected ADT expression (Extended Data Fig. 2b). MMoCHi classification remained accurate when downsampling GEX and/or ADT reads, or when subsampling the number of monocytes or CD8^+^ T_CM_ used for training revealing robustness to lower data quality or fewer, imbalanced, training events (Extended Data Fig. 3). Although this classification uses both ADT and GEX features, MMoCHi was likely insensitive to GEX coverage because both the reference dataset and hierarchy for classification were defined by protein expression.

To compare performance between various methods, we calculated precision, recall and F1 score for each subset, as well as overall accuracy (Fig. 2e, Supplementary Table 5). MMoCHi accurately classified all subsets, with an average F1 score of 0.93 (Fig. 2e: MMoCHi). We tested variations to validate MMoCHi’s design. Performance worsened when training random forests using only GEX and/or when classifying all subsets with only one random forest (instead of a hierarchy) (Fig. 2e: MMoCHi GEX, MMoCHi Flat, and MMoCHi GEX Flat). We noted a modest decrease in performance when relying exclusively on automatic thresholds (Supplementary Fig. 1a,c), and thus encourage users to manually evaluate and adjust automatic thresholds (similar to best-practices for flow cytometry gating^36^). For Fig. 2, we applied MMoCHi using manually defined hyperparameters and the in-danger-noise algorithm enabled (see Methods), but we noted similar performance with MMoCHi’s built-in automatic hyperparameter optimization or in-danger-noise disabled (Supplementary Fig. 1b-c). Overall, these results support MMoCHi’s multimodal, hierarchical, and semi-automated design.

We next compared MMoCHi classification to manual annotation of unsupervised clusters derived from GEX, ADT expression, or the totalVI latent space (see Methods; Fig. 2e: GEX Leiden, ADT Leiden, totalVI Leiden; Extended Data Fig. 2c-g). MMoCHi outperformed all manual annotation methods, particularly GEX Leiden, which failed to identify CD4^+^ T_CM_ or CD8^+^ Naive T cells (Extended Data Fig. 2f). To ensure manual annotation was not hampered by lack of clustering resolution, we also over-clustered the data (see Methods), but this resulted in only a slight improvement (Fig. 2e: GEX Leiden OC, ADT Leiden OC, totalVI Leiden OC). Furthermore, since probabilistic generative modeling on small datasets can be sensitive to feature numbers^37^, we also showed that manual annotation using totalVI was insensitive to the number of input genes (Extended Data Fig. 4a-d).

Lastly, we tested four supervised classification methods for scRNA-seq: ImmClassifier^18^, CellTypist^17^, HieRFIT^19^, and Garnett^20^. We applied ImmClassifier, CellTypist, and HieRFIT as intended, using their pre-designed reference-based training (“pre-trained models”, see Methods). CellTypist was applied to annotate either individual events or by majority voting across clusters (MV), as advised^17^. Garnett was designed, similar to MMoCHi, for classification using marker-based hierarchies,^20^ and was applied using both a published hierarchy, and a custom hierarchy manually selected markers (“Provided Markers” and “Manual Markers” respectively; see Methods; Supplementary Table 6). These pre-trained models include classification of some non-terminal or other, potentially valid, subsets (e.g. T cells, MAIT cells, T_reg_, exhausted T cells; “Alternative”) and classification of populations excluded by our FACS gating (e.g. neutrophils, B cell lineage, NK cells, or γδ T cells; “Erroneous”). Additionally, HieRFIT and Garnett avoid low-confidence classifications, leaving some events undetermined (“Unlabeled”). For benchmarking, we examined only relevantly classified events (i.e. belonging to one of the sorted subsets), but notably, large portions of CellTypist and ImmClassifier classifications were alternative or incorrect, and most Garnett annotations were undetermined (Fig. 2e). Focusing on relevant classifications, all scRNA-seq classifiers performed on-par with GEX Leiden, significantly underperforming compared to MMoCHi classification (Fig. 2e). HieRFIT Pre-trained and Garnett Provided Markers were excluded from benchmarking analyses, as they could not annotate T cells at an appropriate granularity.

All of the supervised approaches described above effectively distinguished monocytes from T cells (Fig. 2e), consistent with previously documented high performance of these tools^17–20^. Our evaluations emphasize the difficulty of segregating T cell subsets that share highly similar transcriptomic profiles without the robust training data and surface protein profiling that MMoCHi leverages. To isolate these two effects from performance of the underlying machine learning algorithms employed, we assessed additional conditions. First, we allowed these tools to “cheat” by supplying them with optimal, batch-free training data—training classifiers using the hashtag-derived sorted subset labels and evaluating performance on a 20% hold-out (“Sort-ref”; see Methods). Although this sort-reference condition is unrealistic (it requires all subsets be FACS-sorted within the same dataset, obviating the need for classification), it revealed differences in performance enabled by MMoCHi’s high-confidence thresholding. Notably, we identified only a modest improvement with idealized training data compared to standard MMoCHi classification, highlighting the efficacy of high-confidence thresholding (Extended Data Fig. 4d). Using sort-reference, classifiers designed for scRNA-seq performed on-par with MMoCHi GEX, but worse than MMoCHi (Extended Data Fig. 4d: comparing within Sort-ref). Although Garnett Sort-ref outperformed MMoCHi GEX Sort-ref, it failed to annotate 20% of the cells (Extended Data Fig. 4d). We next coerced these scRNA-seq classifiers to accept multimodal data (“Multi”; see Methods). Two variants (HieRFIT Multi Sort-ref and Garnett Multi Sort-ref) performed on-par with MMoCHi Sort-ref, suggesting MMoCHi’s underlying machine learning algorithms perform as well as these established tools. Importantly, however, MMoCHi trained using high-confidence thresholds was the only method that could approach this ultra-high accuracy without requiring access to the subset labels from FACS (i.e. “cheating”). Together, these data provide strong support for MMoCHi’s high-confidence thresholding and multimodal training, which together enable effective annotation of these highly similar cell types.

As described above, MMoCHi also has a built-in batch-correction tool for CITE-seq (landmark registration), which we tested on publicly available datasets using diverse antibody panels, chemistries, and sequencing depth (Extended Data Fig. 5a). Landmark registration (run here with no manual input; see Methods) integrated ADT expression more effectively than other ADT normalization techniques (CLR^5^ and dsb^38^), was robust to cell type subsampling, and resulted in a modest improvement in classification performance, especially when the same high-confidence thresholds were applied across batches (Extended Data Fig. 5b-g; Supplementary Table 7).

### MMoCHi integrates classification across diverse human tissue immune cells

We next applied MMoCHi to total immune cells from lymphoid and mucosal tissue samples obtained from human organ donors^17^ (Fig. 3a; Supplementary Table 1). Immune cells were enriched from eight sites across two donors and included lung (LNG), bronchial alveolar lavage (BAL), lung-associated lymph node (LLN), spleen (SPL), jejunum epithelial layer (JEL), jejunum lamina propria (JLP), bone marrow (BOM), and blood (BLD), using methods optimized for each site^17,31^. We performed CITE-seq to profile over 270 surface markers (Supplementary Table 2). In previous analysis of single-cell transcriptomes from this dataset, we detected all lineages of immune cells, including T cells, B cells, innate lymphocytes, and myeloid cells across multiple sites^17^. However, refined immune cell subsets and differences across tissue sites were difficult to resolve in this complex dataset and required extensive manual annotation. Here, we integrated protein and transcriptome profiling using MMoCHi to determine whether transcriptionally similar immune subsets could be identified across diverse tissues.

**Figure 3.**
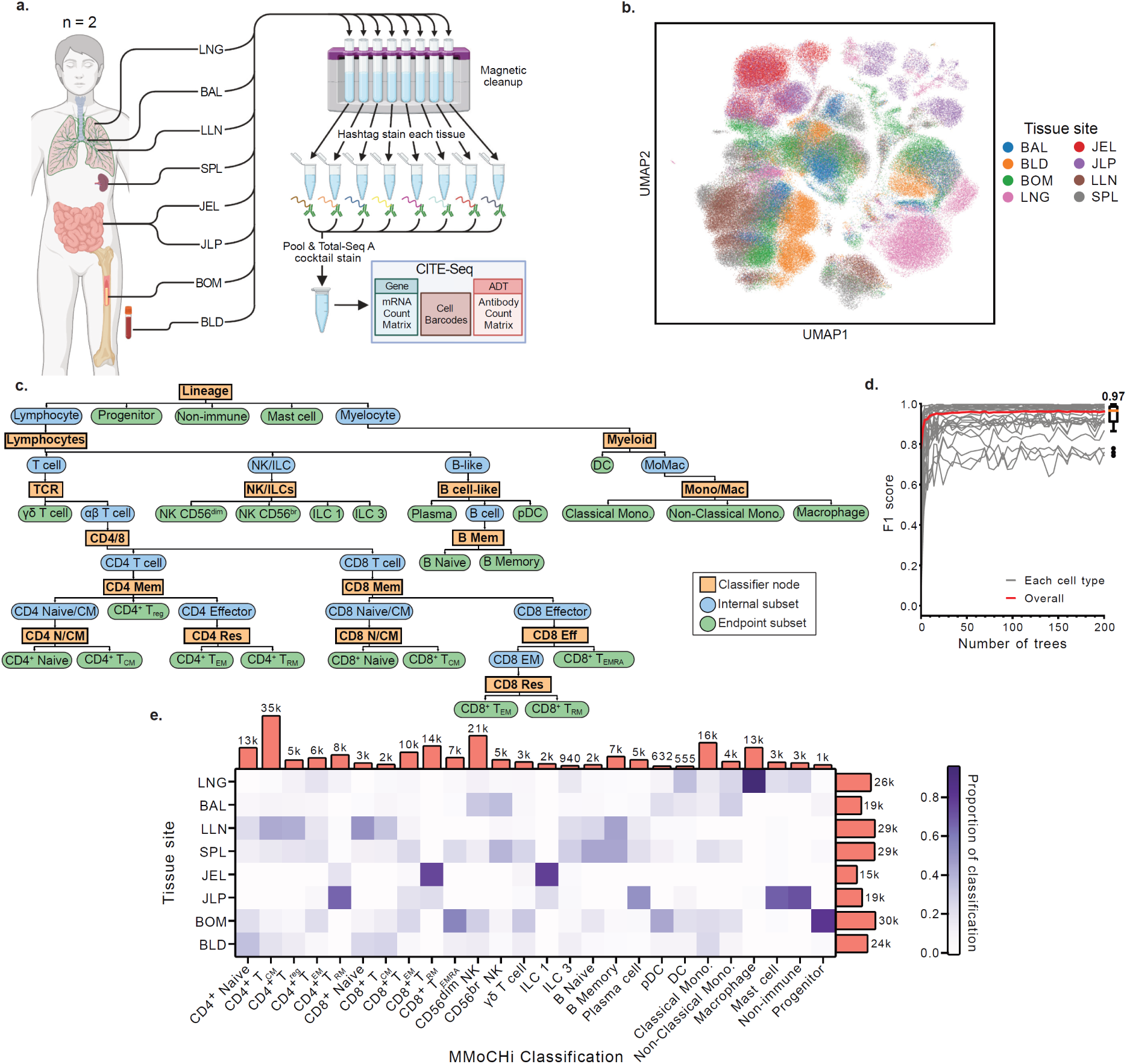
MMoCHi classification applied to human immune cells from blood and diverse tissue sites. a. Single cell suspensions of immune cells were isolated from blood and the indicated tissue sites of two organ donors. CD45^+^ immune cells selected for by magnetic enrichment, hashtagged by tissue site, pooled, and sequenced. b. UMAP of donor-integrated totalVI latent space, colored by tissue site c. MMoCHi hierarchy used for classification (see Supplementary Table 8 for full specification). d. Performance curves showing the F1 score for each endpoint subset when training random forests with various numbers of trees (estimators). F1 scores were calculated on held-out data using high-confidence thresholded events as truth. The red line indicates the median F1 score across all predicted cell types. e. Column-normalized heatmap depicting the distribution of classified cell types across tissue sites. The number of total events in each classification or tissue site are displayed. LNG, lung; BAL, bronchoalveolar lavage; LLN, lung lymph node; SPL, spleen; JEL, jejunum epithelial layer; JLP, jejunum lamina propria; BOM, bone marrow; BLD, blood; T_CM_, central memory T cell; Treg, regulatory T cell; T_EM_, effector memory T cell; T_RM_, resident memory T cell; T_EMRA_, terminally differentiated effector memory T cell; NK, natural killer cell; ILC, innate lymphoid cell; DC, dendritic cell; pDC, plasmacytoid dendritic cell; Mono, monocyte; Mem, Memory; N/CM, Naive/Central Memory; Res, Residency; Eff, Effector. Schematic in (a) created with BioRender.com

For visualization, we embedded a donor-integrated totalVI latent space with UMAP, revealing multiple groupings, some of which corresponded to different tissue sites (Fig. 3b; see Methods). We constructed and applied a MMoCHi hierarchy representing all expected cell types using a combination of transcript and surface protein markers (Fig. 3c; Supplementary Tables 8,9). Training and classifying these approximately 198k immune cell events into 26 subsets took less than 15 minutes (Extended Data Fig. 4, see Methods), demonstrating that MMoCHi’s scalability. The resultant model, using 200 estimators in each random forest, was well-fit at every level of the hierarchy, as measured by the prediction accuracy for internally held-out subsets in the same classification layer and across the full hierarchy (Fig. 3d; Extended Data Fig. 6; Supplementary Fig. 3; see Methods). MMoCHi classified cell types across tissue sites and consistently across donors (Fig. 3e, Extended Data Fig. 7).

We compared MMoCHi classification to manually annotated clusters of the multimodal totalVI latent space (Fig. 4a,b; Supplementary Fig. 4a,b). The methods were broadly concordant, with 72% of events labeled identically; however, disagreements occurred between cell types with similar expression profiles. These disagreements were not resolved by increasing the number of clusters, nor by altering the number of genes input to totalVI (Supplementary Fig. 4c-d). Notably, unsupervised clustering failed to resolve any CD8^+^ T_CM_, γδ T cells, naive B cells, or dendritic cells (DCs). We investigated discrepancies using marker expression (Fig. 4c-e, Supplementary Fig. 5). A substantial percentage (6%) of αβ T cells had conflicting CD4^+^ or CD8^+^ annotations, and by protein and transcript expression MMoCHi classifications were more appropriate (Fig. 4c). Cytotoxic lymphocytes are found across multiple lineages (NK cells, ILCs, CD8^+^ T_EMRA_, and γδ T cells) and are difficult to resolve; however, MMoCHi correctly classified NK cells and ILCs by their lack of CD3 or TCRαβ surface expression, despite expression of *CD3E* and *TRDC* transcripts^24,32^ and their co-clustering with CD8^+^ T_EMRA_ (Fig. 4a,d). MMoCHi also correctly distinguished CD8^+^ T_EMRA_ and γδ T cells where cluster-based manual annotation failed, marked by expression of either CD3 and TCRαβ or CD3, TCRγδ, and variable expression of *TRDV1* and TCRVδ2, respectively (Fig. 4d). MMoCHi also improved identification of T cell memory subsets by expression of CD62L, CCR7, CD45RA, and CD45RO (Fig. 4e). Together, MMoCHi classifications improved identification of known immune cell subsets, particularly in cases of discordant mRNA and protein expression.

**Figure 4.**
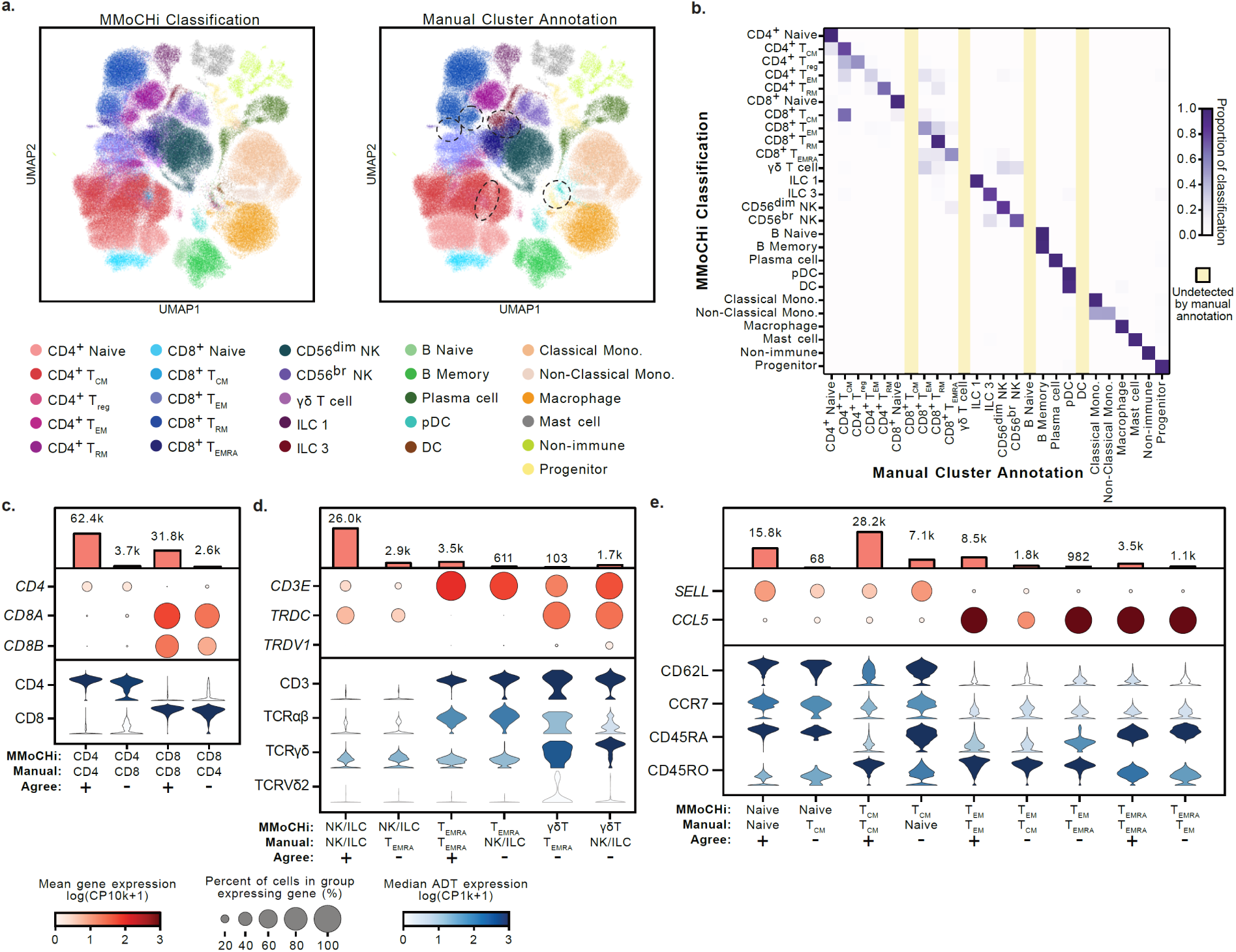
MMoCHi classification of human immune cells outperforms manual annotation by clustering. a. UMAPs of organ donor cells, colored by MMoCHi classification and manual cluster annotations. Dashed ellipses highlight areas of major disagreement between MMoCHi classification and manual annotation. b. Row-normalized heatmap comparing MMoCHi classification to manual annotation. Color represents proportion of cells in each MMoCHi classified population that were manually annotated as each subset. Yellow columns indicate subsets that were not detected by manual annotation. c-e. Plots depicting expression of selected cell type markers on cells grouped by their MMoCHi classification and manual annotation. Dot plots display gene expression (GEX). Dot size represents the percent of cells in the group expressing a gene, and dots are colored by the mean log-normalized GEX counts per ten thousand. Violin plots display the distribution of antibody derived tag (ADT) expression for each population. Violins are colored by the median log-normalized ADT counts per thousand. The number of events in each grouping are displayed above the dot plots. Events are denoted by a “+” where MMoCHi and manual annotation agree, and a “-” where they disagree. T_CM_, central memory T cell; T_reg_, regulatory T cell; T_EM_, effector memory T cell; T_RM_, resident memory T cell; T_EMRA_, terminally differentiated effector memory T cell; ILC, innate lymphoid cell; NK, natural killer cell; pDC, plasmacytoid dendritic cell; DC, dendritic cell; Mono, monocyte

We additionally sought to evaluate MMoCHi’s performance when pre-trained classifiers are applied to new datasets. To this end, we applied MMoCHi classifiers trained using the FACS dataset (“FACS-trained”) to the αβ T cells and monocytes isolated from the PBMCs of organ donors (Extended Data Fig. 8a-b; see Methods). The FACS-trained MMoCHi model outperformed pre-trained or FACS-trained ImmClassifier, CellTypist, or Garnett models at replicating the correct annotation for each cell type (Extended Data Fig. 8a-f, Supplementary Table 10). Although we do not anticipate that this will be a primary use-case of the algorithm, these data further support MMoCHi’s efficacy.

### MMoCHi classifiers are highly interpretable and identify cell type markers

Having demonstrated MMoCHi’s utility for cell type annotation, we next determined which features (transcript and surface protein expression) were useful for subset delineation. Many single-cell classifiers are trained on dimensionally reduced data^21^, hampering feature-level interpretability, but random forests within MMoCHi are trained using all available protein-coding genes and surface proteins. During training, features are selected for their contribution to decreasing impurity, thus providing a natural ranking of features by their importance for classification^39,40^ (see Methods). We analyzed the important features at each level of the hierarchy, which included subset markers defined by the user (for high-confidence thresholding) and revealed other known subset markers, identified *de novo*, including: *IL1R1* and *GATA2* for mast cells^41^, *APOE*, *ACP5*, and *C1QC* for macrophages^42^, *CD79A*, *MARCKS*, *MEF2C*, and CD32 for B cell-like^43–46^, and *TYROBP*, *GZMB*, and CD123 for pDCs^47,48^ (Extended Data Fig. 9; Supplementary Tables 8,11-12). This demonstrates that MMoCHi learns informative cell type representations— an important pre-requisite for novel marker identification.

Next, we wondered if MMoCHi could be leveraged to improve transcriptome-based segregation of naive T cells and T_CM_, which are defined by specific CD45 isoform surface-expression but share similar transcriptomets^1,18,28,30^. First, we trained GEX classifiers using high-confidence CD4^+^ and CD8^+^ Naive/T_CM_ selected by multimodal thresholding (Supplementary Table 8). Despite excluding ADTs from training, these classifiers were well-fit, highly accurate, and resulting subsets had expected surface marker expression (Fig. 5a,b). To identify GEX markers of these subsets, we next interrogated impurity-based important features (Fig. 5c,d; Supplementary Tables 13-14). We applied robust filtering to highlight genes with clear differences in expression and reduce the potential for tissue-specific contamination (see Methods), and identified a suite of memory-associated genes, including *ITGB1*, *LMNA*, and *FAM129A*^27,30^ as markers upregulated in CD4^+^ and CD8^+^ T_CM_. Improved prediction accuracy by classifiers trained with only the top 1000 important features confirmed that these features were useful for GEX classification (Fig. 5e).

**Figure 5.**
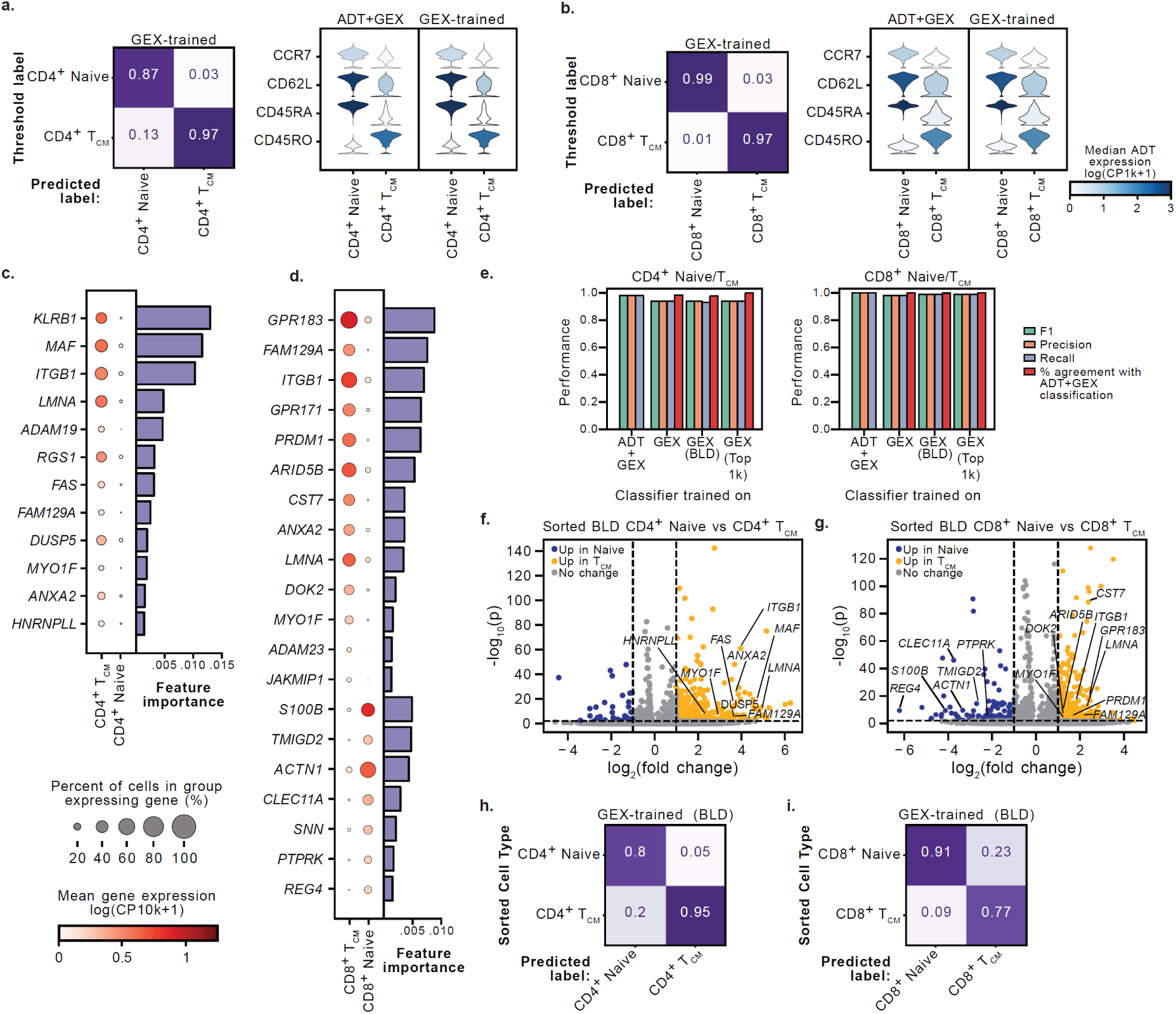
MMoCHi reveals additional transcript markers distinguishing naive and central memory T cells. a-b. Performance of gene expression (GEX)-only classification of Naive/T_CM_ CD4^+^ (a) and CD8^+^ (b) T cells. Column-normalized confusion matrices (left) comparing classification to held-out, high-confidence threshold labels. Violin plots (right) display the distribution of antibody derived tag (ADT) expression for each population as classified by MMoCHi trained on multimodal data (ADT+GEX) or GEX-only. Violins are colored by the median log-normalized ADT counts per thousand. c-d. Dot plots displaying expression of important features for GEX-only classification of Naive/T_CM_ CD4^+^ (c) or CD8^+^ (d) T cells. Dot size represents the percent of cells in the group expressing a gene, and dots are colored by the mean log-normalized GEX counts per ten thousand. The top important features associated with a single subset (log_2_(fold change) > 2 and greater than 10% change in dropout rate) are displayed. All features displayed were within the top 1000 important features when training with GEX-only on all tissues, and with blood (BLD) only. The importance of each feature is shown in a bar chart (right). All displayed features were significantly differentially expressed (p < 0.05). e. Bar plots comparing performance of Naive/T_CM_ CD4^+^ and CD8^+^ T cell classifiers trained using multimodal data (ADT+GEX), GEX only, GEX with only BLD cells, or only the top 1000 important GEX features. f-g. Volcano plots displaying differential gene expression between sorted blood Naive and T_CM_ CD4^+^ (f) or CD8^+^ (g) T cells. Points represent genes and are colored by differential expression (|log_2_(fold change)| > 1 and P < 0.05). Important features for classification that were differentially expressed are highlighted. h-i. Column-normalized heatmaps displaying performance of GEX only classifiers trained on organ donor BLD applied to sorted Naive/T_CM_ CD4^+^ (h) and CD8^+^ (i) T cells. Statistical significance was calculated using a two-sided Wilcoxon with tie correction, followed by a Benjamini–Hochberg adjustment for multiple comparisons. T_CM_, central memory T cell.

We sought to validate these findings using our scRNA-seq of sorted CD4^+^ and CD8^+^ naive and T_CM_ populations. Most markers identified for naive and T_CM_ were also differentially expressed within the sorted dataset (Fig. 5f,g; Supplementary Table 15). Random forests trained on organ donor PBMCs were well-fit, effectively recapitulating the sorted labels (Fig. 5h,i). Overall, these findings demonstrate the capacity of MMoCHi to leverage CITE-seq to identify gene expression markers and train classifiers that can be effectively applied to scRNA-seq datasets.

### MMoCHi effectively annotates other multimodal datasets and cell types

Lastly, we explored whether MMoCHi could be easily extended beyond CITE-seq. First, we applied MMoCHi to publicly available paired transcriptome and surface proteome of sorted T and NK cells generated using the Ab-seq platform^7,49^. We obtained high concordance with cluster-based annotations and expected marker expression (Fig. 6a,b; Extended Data Fig. 10a; Supplementary Fig. 6a,d; Supplementary Tables 16-17). Next, we applied MMoCHi to scRNA-seq of a high-grade glioma biopsy^50^. In these data, malignant cells were challenging to identify based on marker genes, because they are highly heterogenous and express many of the same genes as non-neoplastic cells in the brain microenvironment. However, gliomas often harbor clonal aneuploidies that are detectable by averaging gene expression across whole-chromosomes^50^. Using a joint feature set comprised of transcriptome and computed whole-chromosome expression (see Methods), MMoCHi effectively segregated tumor cells from other non-neoplastic subsets, which MMoCHi also annotated accurately (>98% agreement overall; Fig. 6c,d; Extended Data Fig. 10b; Supplementary Fig. 6b,e; Supplementary Tables 16-17). Notably, chromosomal expression was useful both as a marker for high-confidence thresholding and important internally for classification (Extended Data Fig. 10b; Supplementary Table 17).

**Figure 6.**
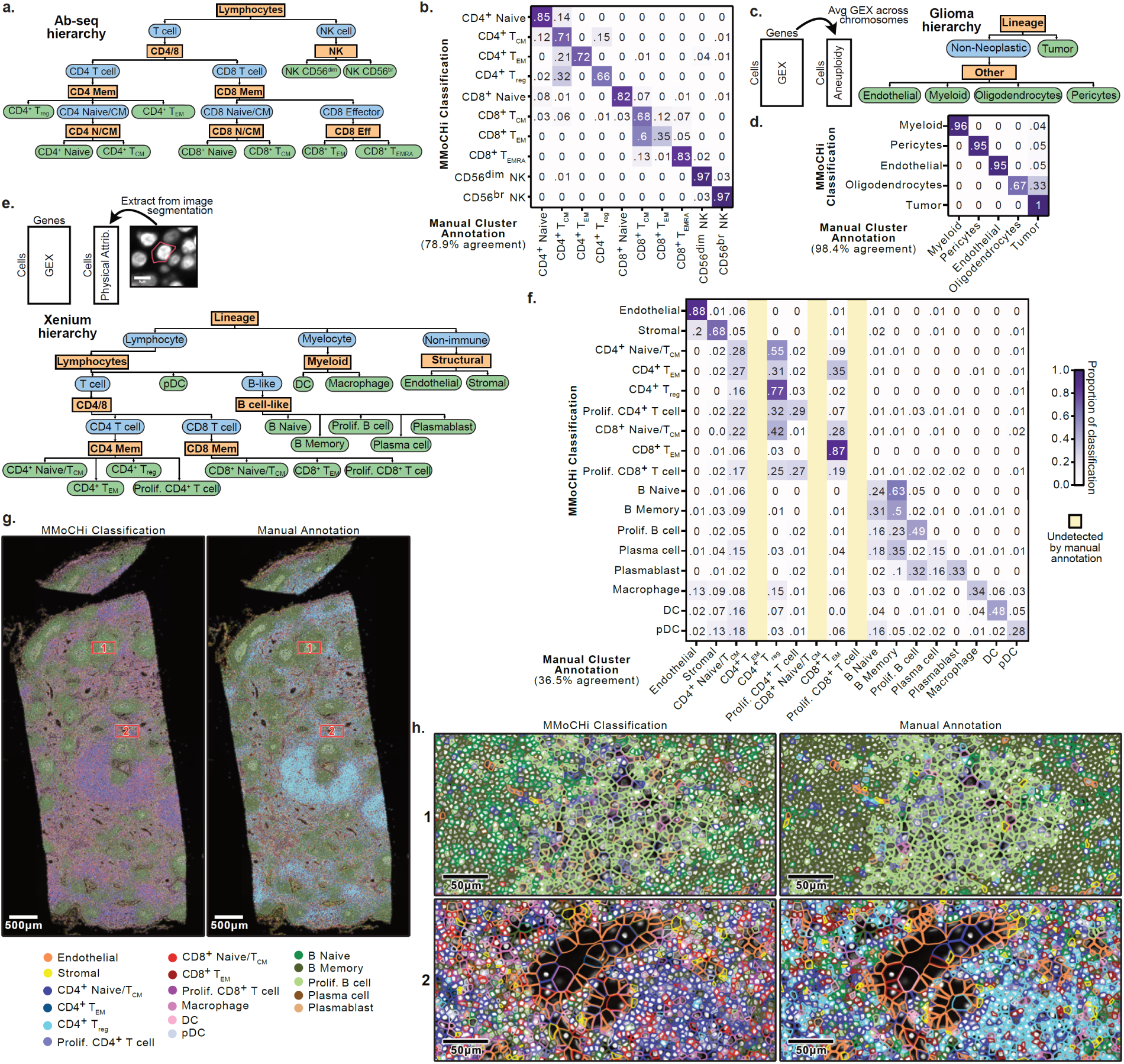
MMoCHi applied to other multimodal datasets. MMoCHi applied to Ab-seq data of sorted T and NK cells (a-b), a scRNA-seq of a high-grade glioma biopsy (c-d), or spatial transcriptomics of a human lymph node using the 10x Xenium platform (e-h). a, c, e. MMoCHi hierarchies used for classification (see Supplementary Table 17 for full specification) and schematics for extraction of multimodal data for classification (see Methods). b, d, f. Row-normalized heatmaps comparing MMoCHi classification to manual annotation. Color represents proportion of cells in each MMoCHi classified population that were manually annotated as each subset. Yellow columns indicate subsets that were not detected by manual annotation. g. Images displaying DAPI staining intensity on a human lymph node, overlaid with detected cell boundaries (see Methods). Cell boundaries are colored by MMoCHi classification or manual annotation. h. Magnified sections of the images in g, as outlined by the red boxes.

We further explored whether MMoCHi to be adapted to spatial profiling, using a human lymph node dataset generated by the Xenium platform (10x Genomics), which enables multiplexed profiling of 100s of RNAs *in situ* with subcellular resolution. By pairing the 377 profiled genes with physical attributes of each cell extracted from the image and segmentation (see Methods), MMoCHi effectively classified 17 subsets, revealing underlying spatial structure, including B cell follicles and lymphatics (Fig. 6e-h; Supplementary Table 16). MMoCHi outperformed manual cluster-based annotation, which failed to segregate CD4^+^/CD8^+^ T cells and various B cell-lineage subsets (Fig. 6f; Extended Data Fig. 10c; Supplementary Fig. 6 c,f). As before, the added modality was useful for MMoCHi classification, where physical attributes (particularly nuclear circularity and DAPI intensity), ranked in the top 5% of features used (Extended Data Fig. 10c; Supplementary Table 17). Overall, MMoCHi’s success in annotating these datasets of varying subset compositions and modalities demonstrates the algorithm’s high adaptability to classification of multimodal single-cell datasets.

## DISCUSSION

The advent of multimodal single-cell technologies has enabled high-dimensional profiling of many systems, organs, diseases, and species. However, the development of analytical tools to identify cell states and their features consistently across multimodal studies is lagging behind these data acquisition technologies. Here, we present MMoCHi, a multimodal, hierarchical classification approach for cell type annotation of CITE-seq data, which integrates both gene and protein expression, does not require reference datasets, performs landmark registration for harmonization across large datasets, and accurately classifies both closely related subsets within a single lineage and diverse subsets from across lineages. We devised an experimental validation approach that allowed us to directly compare MMoCHi and several established annotation modalities directly to FACS-based subsetting for the exact same sample of cells. This enabled an extensive benchmarking study across conventional clustering-based methods, supervised machine learning algorithms, and both reference-based and marker-based classifiers with a uniform and independent reference annotation. Applied to immune cell CITE-seq datasets, MMoCHi outperforms current annotation algorithms, and identifies new markers for subset delineation. Together, MMoCHi provides an adaptable approach for applying marker-based annotation to multimodal datasets.

Currently available single-cell classifiers are primarily designed for scRNA-seq; however, CITE-seq can be leveraged to improve annotation^5,6^. MMoCHi robustly classifies closely related immune cell subsets with discordant transcriptome and surface proteome profiles. Here we show that MMoCHi effectively distinguishes T cell subsets defined by surface expression of CD45 isoforms and homing receptors but share overlapping transcriptomes^1,18,28,30^—namely naive T cells from T_CM_ and T_EM_ from T_EMRA_. By classifying hierarchically, MMoCHi was also able to correctly annotate diverse immune lineages without compromising its performance at segregating functionally and transcriptionally similar cell types, including cytotoxic NK cells, CD8+ T_EMRA_, and γδ T cells^24,32,33,51^. MMoCHi can accurately classify cell types despite other sources of variation, as shown by integrated classification of multiple immune cell lineages across blood and 8 disparate tissue sites of two organ donors.

By providing a platform for using high-confidence, marker-based thresholding for selection of training data, MMoCHi enables the flexible design of new classifiers based on prior knowledge without a requirement for carefully curated reference atlases. In particular, applying high-confidence thresholding to CITE-seq protein expression serves as a natural extension of the vast knowledge-base developed of flow cytometry markers and gating^26,36^. Similar to other marker-based strategies, these subset definitions are highly interpretable and can be robustly applied across studies and specific sequencing conditions to train new classifiers^20,25^. To reduce redundancy in efforts to design MMoCHi hierarchies, the hierarchies defined in this manuscript, along with other community contributions, have been made available online (https://mmochi.readthedocs.io). Here, we also apply MMoCHi’s multimodal training data selection to improve transcriptome-based annotation of T cell subsets across datasets, demonstrating potential for MMoCHi to advance annotation of scRNA-seq data as well.

In contrast with efforts to automate cell type annotation^16,17^, MMoCHi’s thresholding schemas require careful marker curation and domain expertise, however; by leveraging this expertise, MMoCHi classifications may better reflect canonically defined, and biologically relevant designations. Additionally, MMoCHi is not optimized for novel subset identification, requiring the user to define all cell types within a sample for classification. Thus, MMoCHi is complemented by multimodal unsupervised exploration of the dataset either before or after classification to annotate or identify markers of unexpected cell types and states, or to identify novel subsets of canonical cell types.

Beyond cell type classification, MMoCHi learns key features of protein and gene expression, which can be used to identify new cell type markers and derive biological insights. MMoCHi random forests do not require prior dimensionality reduction, enabling evaluation of individual surface proteins and protein-coding genes as subset markers. In classifying diverse and highly similar subsets, we show that MMoCHi extracts relevant markers, relying on both features used for training data selection, and other known gene and protein markers of each subset. In using MMoCHi to identify improved transcriptional markers of naive and central memory (T_CM_) T cells, we discovered that circulating T_CM_ actually upregulate markers that had previously been associated with T cells in tissue^30,31,52^ and effector memory T cells^27,30^. This finding raises the possibility that early stages of tissue programming occur during T_CM_ differentiation and indicate a gradual transition between T cell memory subsets. Importantly, we did not previously identify these expression patterns of circulating T_CM_ in earlier studies of T cells in tissues and blood^30,31^, likely due to the lack of CITE-seq and multimodal classification.

While we have developed MMoCHi with CITE-seq applications in mind, the algorithm is designed for easy extension to other modalities frequently paired with single-cell transcriptomes, such as profiling chromatin accessibility, T and B cell receptors, mutational landscapes, intracellular proteins, or a combination of these modalities^53–58^. We also anticipate applications to emerging technologies for multimodal, single-cell spatial profiling^59,60^. We demonstrated the potential generality of MMoCHi through extension to joint aneuploidy/gene expression profiles from a solid tumor and joint classification of *in situ* spatial profiling data using morphological and gene expression features measured using the 10x Genomics Xenium platform. While we mainly focused on immunology, cell type classification is a ubiquitous problem in single-cell genomics. Thus, we expect broad utility for MMoCHi in diverse biological applications, including identification of developmental states, building atlases of complex tissues and tumors, profiling model organisms, and analyzing clinical specimens.

## ACKNOWLEDGEMENTS

We thank Joshua I. Gray and Rory E. Morrison-Colvin for helpful discussions and members of the Farber laboratory for help with tissue processing. This work was supported by a Seed Networks for the Human Cell Atlas grant from the Chan Zuckerberg Initiative (CZF2019-002452) and NIH grants AI128949 and AI106697 awarded to P.A.Si. and D.L.F. D.P.C. was supported by the Columbia University Graduate Training Program in Microbiology and Immunology (T32AI106711). P.A.Sz. was supported by a Canadian Institutes of Health Research (CIHR) Fellowship. Research reported here was performed in the Columbia Stem Cell Initiative Flow Cytometry Core, the Sulzberger Columbia Genome Center, and the Columbia Single Cell Analysis Core (supported by grant P30CA013696).

The content is solely the responsibility of the authors and does not necessarily represent the official views of the NIH. We wish to thank the donor families for their generosity and the exceptional efforts of the transplant coordinators and staff of LiveOnNY for making this study possible.

## AUTHOR CONTRIBUTIONS

D.P.C. designed and performed experiments, analyzed data, made figures, and wrote the manuscript. W.L.S. performed experiments, analyzed data, made figures, and edited the manuscript. D.C., S.B.W., P.A.Sz., and I.J.J. prepared samples for single cell sequencing. P.A.Si. and D.L.F. designed experiments, analyzed data, wrote, and edited the manuscript.

## METHODS

### MMoCHi

MMoCHi is designed to standardize classification of cell subsets using multimodal (e.g. CITE-seq) data (Fig. 1). To train a new MMoCHi model, the user specifies a hierarchy of cell subsets, along with marker definitions (based on thresholds of either gene or protein expression) for each. Similar to other hierarchical classifiers,^19,20^ hierarchies progressively segregate subsets by similarity of expression profiles, which may or may not reflect a cell type’s ontogeny. MMoCHi iterates through each level of the hierarchy, selecting high-confidence members of each subset, training a random forest classifier on normalized gene and protein expression, and classifying cell types. Once trained, MMoCHi classifiers can be interrogated for feature importances or applied to extend cell type annotation to other datasets.

### Feature selection, normalization, and batch correction

As input to MMoCHi, gene expression (GEX) and antibody derived tag (ADT) expression were log-normalized. For many markers, ADT expression distributions matched the expected bimodal, trimodal, or gradients of expression observed by flow cytometry. To account for variation in antibody staining across samples, ADT expression in multi-sample datasets was batch-corrected using a landmark registration algorithm (Fig. 1a), similar to previously described techniques for flow cytometry and CITE-seq^34,35^. Landmark registration was applied to expression of all ADTs, except for isotype controls, which were excluded from downstream analyses. First, landmarks (peaks) were identified in the distribution of expression for each ADT in each sample as automatically detected local maxima (*scipy.signal.find_peaks*, using the Scipy package v1.10.1) on kernel-density-smoothed (*scipy.stats.gaussian_kde*) ADT expression, or manually identified. Curve registration and warping functions were applied to align these landmarks across samples (*skfda.preprocessing.registration.landmark_elastic_registration_warping*, using the scikit-fda package v0.8.1). For each ADT in each sample, the lowest detected peak (assumed to represent “negative” expression) and the uppermost peak (assumed to represent “positive” expression) were aligned to center on fixed values: 1 and 3, respectively. In samples with only one peak detected, the peak was aligned to the “negative” location, although this could be adjusted manually to align batches with only positive-expression for an ADT. As a batch-correction technique, landmark registration assumes that changes across samples in the degree of marker positivity on marker-positive populations is due to technical artifacts, not biological differences. This is a reliable assumption for annotation purposes, especially when samples are from similar sources or contain the same cell types at varying frequencies.^34^ Once batch-corrected, thresholds delineating positive and negative populations for ADT markers could be applied across batches.

### Selection of training data

At each level of the hierarchy, we selected high-confidence events for each subset using a combination of automatically-derived and manually-adjusted thresholds on the user-supplied marker definitions (Fig. 1b, Supplementary Tables 4,7,8,16). ADT markers were fitted with 1- or 2- component Gaussian mixture models (*sklearn.mixture.GaussianMixture*, using the Sklearn package v1.3.0), and thresholds were automatically defined as 1 to 4 standard deviations above or below the means of each Gaussian. ADT thresholds were then manually refined using established flow cytometry gating principles (e.g. Ref.^36^). GEX thresholds were drawn similar to Garnett^20^, primarily capturing events with any marker expression, or occasionally (in genes with high background expression) capturing only the highest expressing events. A portion (20%) of these high-confidence events were held out for testing and validation, and the remaining 80% could be used for training. We then resampled this training dataset (Extended Data Fig. 1a) to remove events likely to be mistakenly labeled due to imperfect marker thresholding (noise) and to overrepresent events likely to be misclassified (in danger). To identify “noise” and “in danger” events, principal component analysis (PCA; *scanpy.pp.pca*, using the Scanpy package v1.9.3) was run on the scaled expression of the top 5000 highly variable genes (*scanpy.pp.highly_variable_genes*) and all ADTs. The 5 nearest neighbors were calculated for each event (*sklearn.neighbors.NearestNeighbors*). Events were considered “noise” and removed if all neighbors disagreed with their high-confidence label. Events were considered “in danger” if less than half the neighbors agreed with their high-confidence label. Training events were also clustered using the Leiden algorithm^61^ (*scanpy.tl.leiden*) to identify “in danger” clusters—clusters representing less than 5% of a subset’s training events. All events considered “in danger” were oversampled 5 times to increase representation during training. At classification nodes where subsets were not expected to segregate by unsupervised approaches—including Lymphocytes and TCR—these selection steps were skipped. To account for class imbalance in the training dataset, events for all subsets were oversampled to equal numbers. Finally, training events were subsampled without replacement to a maximum of 20,000 events for computational performance. The selection of training data occurred separately for each batch. If a minimum of 100 high-confidence events were not identified in a batch, events would be spiked into that batch from other batches (Extended Data Fig. 1b).

### Training and calibration of random forest classifiers

At each level of the hierarchy, we trained a random forest (*sklearn.ensemble. RandomForestClassifier*) with a default of 100 trees, each with a max depth of 25 to reduce overfitting. In datasets with multiple batches, 100 trees were trained separately for each batch and added to the forest. Once trained, the forest would predict the subset identity of events and provide the proportion of trees in agreement with the classification. To convert these proportions of trees in agreement to probabilities, they were transformed using an isotonic regression (*sklearn.calibration.CalibratedClassifierCV*) trained on a subset of the held-out data^62^. Classification performance was then evaluated on the remaining hold-out data.

Hyperparameters were tuned in cases of poor fit, either automatically or manually. Hyperparameter tuning on the underlying random forest classifiers focused on optimizing values for the number of trees in the forest (*n_estimators*), the maximum depth of each tree (*max_depth*), and the maximum number of features to consider when looking for the best split during random forest training (*max_features*). For each hyperparameter tested, fit was determined by calculating the balanced accuracy score (*sklearn.metrics.balanced_accuracy_score*) between predicted and high-confidence labels on a subset of the held-out data (distinct from the subset used for performance evaluation). For automatic optimization, *n_estimators* were tested at 50, 100 (MMoCHi default), 200, 400, 800, and 1200; *max_depth* of 10, 25 (MMoCHi default), or unlimited; and *max_features* of log_2_(total), square-root of total (MMoCHi default), 5% of total, or 10% of total features. Optimal hyperparameter values were selected as either the maximum score, or as the lowest value before this maximum score where score did not improve by at least 0.004. To increase fit, the *max_features* hyperparameter was manually set to 10% of total features for CD4^+^ and CD8^+^ memory classification for sorted T cells, as well as Lymphocyte, TCR, and CD4/8 classification for organ donor cells.

The trained classifier was then used to predict subset identity for all events at that level. This process of training data selection, classifier training, and prediction were repeated until all cells had been labeled to terminal subsets.

### CITE-seq profiling of FACS sorted subsets

Peripheral blood from a consenting healthy volunteer (34-year-old male; Supplementary Table 1) was obtained by venous puncture, through a protocol approved by the Columbia University IRB and complying with relevant ethical regulations for work with human participants.

Peripheral blood mononuclear cells (PBMCs) were isolated using RosetteSep Granulocyte Depletion Cocktail (StemCell Technologies), following manufacturer’s protocols for density gradient centrifugation. Briefly, samples were incubated with the cocktail at 20°C for 10 minutes, diluted 1:1 with FACS buffer (DPBS 10% FBS 2mM EDTA), then layered over Ficoll-Plaque in SepMate PBMC isolation tubes (StemCell Technologies). Samples were centrifuged at 1200 x g for 10 minutes at 20°C, and PBMC layers were isolated according to instructions. Samples were washed (400 x g for 10 minutes) with FACS buffer. For further erythrocyte removal, pellets were resuspended in ACK lysis buffer (Gibco) incubated for 2 minutes at 37°C and washed with FACS buffer.

PBMCs were stained with Zombie NIR Fixable Viability dye (BioLegend) for 30 minutes. Samples were kept at 4°C in the dark. Cells were washed thrice with FACS buffer, resuspended in TrueStain FcX and TrueStain Monocyte Blocker (BioLegend), and incubated for 10 minutes. We designed a FACS-sort antibody cocktail, prioritizing antibody clones with discrete epitopes from the TotalSeq-A Universal Human Panel (BioLegend) to reduce steric hinderance during CITE-seq staining (Supplementary Table 2). Cells were incubated with the FACS-sort antibody cocktail for 30 minutes, then washed thrice with FACS buffer. Cells were incubated in TrueStain FcX (BioLegend) for 10 minutes, then stained using a custom TotalSeq-A Universal Human Panel (BioLegend) for 30 minutes, according to manufacturer instructions. Samples were washed thrice with FACS buffer. Cells were sorted using a FACS Aria II (BD Biosciences; Extended Data Fig. 2a). Seven T cell memory populations (CD4^+^ Naive, CD4^+^ T_CM_, CD4^+^ T_EM_, CD8^+^ Naive, CD8^+^ T_CM_, CD8^+^ T_EM_, CD8^+^ T_EMRA_), and monocytes were sorted into sterile, heat inactivated FBS. Sort purity was calculated as the number of events falling within a subset’s gates divided by the total number of singlet events times 100, using the same gating strategy as the sort (Extended Data Fig. 2a). All sorts were high purity, with mean purity 94.4% (Supplementary Table 3). Sorted populations were washed in FACS buffer and resuspended in TrueStain FcX (BioLegend) for 10 minutes. Samples were then stained with TotalSeq-A hashtag antibodies (BioLegend). In this experiment, the hashtag-oligos (HTOs) correspond to the sorted immune cell subsets (HTO1: CD4^+^ Naive, HTO2: CD4^+^ T_CM_, HTO3: CD4^+^ T_EM_, HTO4: CD8^+^ Naive, HTO5: CD8^+^ T_CM_, HTO6: CD8^+^ T_EM_, HTO7: CD8^+^ T_EMRA_, HTO8: Monocytes). Samples were then washed thrice with FACS buffer, and pooled.

### CITE-seq profiling of human tissue samples

Human tissues were obtained from deceased organ donors as previously described^17,31,63,64^. The use of tissues from organ donors is not considered human subjects research as confirmed by the Columbia University IRB because the donors are deceased. Mononuclear cells (MNCs) were isolated from tissue sites of two organ donors (D496 and D503; Supplementary Table 1), as described^17^. Approximately 1 million MNCs per tissue site were washed in FACS buffer and resuspended in TrueStain FcX (BioLegend) for 10 minutes. Samples were then stained with TotalSeq-A hashtag antibodies (BioLegend) for each tissue. For each donor, the hash-tagged MNCs from each tissue site were pooled, washed with FACS buffer, and stained with the TotalSeq-A Human Universal Cocktail panel according to the manufacturer’s instructions (BioLegend).

### Library preparation, sequencing, and alignment

The sorted immune cells were counted, diluted to an appropriate volume, and loaded across two lanes of a 10X Genomics Chromium instrument targeting 6,000 cells each. Samples from each organ donor were loaded across 16 lanes of a 10X Genomics Chromium instrument targeting 10,000 cells each. cDNA synthesis, amplification and sequencing libraries were generated using the Next GEM Single Cell 3’ Kit v3.1 (10X Genomics) with the recommended modifications for compatibility with the TotalSeq-A cell hashing and CITE-seq reagents (BioLegend). Organ donor GEX libraries were sequenced on a NovaSeq 6000 (Illumina) with 100 cycles for reads 1 and 2. Organ donor ADT and HTO libraries were sequenced on a NextSeq 500 (Illumina) with 28 cycles for read 1 and 55 cycles for read 2. All libraries for FACS sorted subsets were sequenced on a NextSeq 500 (Illumina) with 28 cycles for read 1 and 44 cycles for read 2.

Reads were analyzed by pseudoalignment using kallisto v0.46.2 (GRCh38 with Gencode v24 annotation) and bustools v0.40.0^65–67^. CITE-seq and hashtag barcodes were demultiplexed and extracted using DropSeqPipeline8, as previously described^68^. Hashtags were demultiplexed by CLR normalization, k-means clustering, and statistical identification of singlets by fitting a negative binomial model as described^5^. For both datasets, the GEX matrices were normalized to log(counts*10,000/total_counts+1), and ADT expression matrix were normalized to log(counts*1,000/total_counts+1).

### Analysis and benchmarking using sorted T cells

Cells from the T cell sort were filtered to remove events with fewer than 1000 unique counts, fewer than 200 genes detected, or over 10% mitochondrial counts. A MMoCHi hierarchy was developed (Fig. 2b) and classification performed using the algorithm above. In flat classification variations (MMoCHi Flat, MMoCHi GEX Flat), high-confidence thresholding and classification were performed for all subsets in a single classification node. In GEX variations (e.g. MMoCHi GEX, MMoCHi GEX Flat, etc.), ADT expression data was excluded from the detection of in-danger noise events and random forest training. In “Sort-ref” variants (e.g. MMoCHi Sort-ref, MMoCHi GEX Sort-ref, etc.), training events were selected using the sort labels instead of high-confidence thresholding. For benchmarking tests, to mirror intra-dataset performance testing of other reference-based tools^17,19^ and avoid leaky pre-processing bias^69^ a portion of the dataset (20%) was set aside to only be used for performance evaluation (completely held-out from the delineation of high-confidence thresholds, training data selection, random forest training, random forest calibration, and any manual or automatic hyperparameter optimization). To test MMoCHi’s robustness, performance was also tested by subsampling training event numbers and downsampling reads. Random subsampling was performed on the number of monocyte or CD8^+^

T_CM_ events included in the training dataset, down to 5% of the total of monocyte/CD8^+^ T_CM_ events profiled. Aligned reads for GEX and ADT libraries were downsampled prior to generation of raw-count expression matrices. Downsampling was performed down to 1% of the total aligned reads, and the effect of this downsampling on total counts is displayed.

We manually annotated unsupervised clusters by average expression of gene and protein markers. Leiden clusters of gene expression (GEX Leiden) were computed on a PCA of the highly variable genes with Scanpy^70^ defaults. Leiden clusters of ADT expression (ADT Leiden) were computed on a PCA of all ADTs except for isotype controls. totalVI latent space was computed on all ADTs except for isotype controls, and highly variable genes selected using top 4000 genes as defined by the Seurat v3 method^71^, as recommended^14^ (*scvi.model.TOTALVI.train* and *scvi.model.TOTALVI.get_latent_representation* using scvi-tools v1.0.2). To ensure results of totalVI were not sensitive to the number of features used, we also computed totalVI latent spaces using the top 1000, 2000, and 8000 highly variable genes. Leiden clustering was performed on the 10 nearest neighbors of the top 40 principal components or the entire totalVI latent space. Over-clustering was also computed where highly variable gene selection, dimensionality reduction, and Leiden clustering were repeated to sub-cluster each cluster (excluding clusters of fewer than 40 events) resulting in 72 GEX clusters (GEX Leiden OC), 128 ADT clusters (ADT Leiden OC), and 116 totalVI clusters (totalVI Leiden OC). For visualization, UMAP (*scanpy.tl.umap*) embeddings of each of these feature spaces were calculated using defaults.

ImmClassifier^18^, CellTypist^17^, and HieRFIT^19^ were applied as recommended, using pre-trained models downloaded from online. ImmClassifier was run using *docker* with the built-in model, trained by inputting a log_2_(counts*10,000/total_counts+1) transformed GEX matrix, as recommended. CellTypist v1.6.2 was run by applying the “Healthy_COVID19_PBMC” model (trained on PBMCs from healthy individuals and individuals with COVID-19^72^) with and without majority voting enabled (*celltypist.annotate*; CellTypist MV and CellTypist, respectively). A HieRFIT v0.1.0 model trained on 10x PBMC data was applied (*HieRFIT::HieRFIT*) with defaults settings.

Garnett^20^ models were trained using high-confidence thresholding with the provided online hierarchy (Provided Markers) and on manually and automatically (*garnett::top_markers*) selected transcript markers which were evaluated using built-in functions (*garnett::check_markers*; Manual Markers; Supplementary Table 6). Garnett v0.2.20 models were trained (*garnett::train_cell_classifier*) with marker propagation disabled and prediction (*garnett::classify_cells)* was performed with cluster extension disabled.

Additionally, CellTypist, HieRFIT, and Garnett were trained using the HTO-derived sort labels as a reference dataset and a evaluated using a 20% hold-out (“Sort-ref”). A CellTypist model was trained (*celltypist.train*) with two-pass training enabled for feature selection. This model was applied to held-out data with and without majority voting enabled (CellTypist MV Sort-ref and CellTypist Sort-ref, respectively). HieRFIT and Garnett models were both trained (*HieRFIT::CreateHieR*; *garnett::train_cell_classifier*) using a hierarchy structured identically to the MMoCHi hierarchy (Fig. 2b) and applied to held-out data (HieRFIT Sort-ref and Garnett Sort-ref, respectively). The three models were also retrained using the combined multimodal matrix of GEX and ADT expression (“Multi”). To coerce these tools to accept the multimodal matrix, the *check_expression* parameter in CellTypist functions and the *db* parameter in Garnett functions were disabled.

Precision, recall, F1 scores and overall accuracy were calculated using sort labels as truth (*sklearn.metrics.precision_recall_fscore_support*; *sklearn.metrics.accuracy_score*). Garnett and HieRFIT can provide unknown and intermediate cell type labels, which were quantified and excluded from performance metrics calculations. Pre-trained classifier models could also provide erroneous (e.g. not included in the FACS sort) or alternative (i.e. potentially valid annotations but not using the subsetting-axis of interest) cell type labels, which were also quantified, then excluded from performance calculations (Supplementary Table).

### Analysis of ADT batch-correction with published CITE-seq data

To test landmark registration against other published CITE-seq normalization techniques, we used four public datasets of CITE-seq performed on healthy PBMCs available from 10x Genomics (Extended Data Fig. 5a). Each dataset was treated as a separate batch. For each batch, log(counts*1,000/total_counts+1) and landmark registration were performed as described above. For these data, no manual modifications were made to the built-in automatic peak detection. CLR and dsb normalizations were performed using the Muon package^73^ v0.1.5 (*muon.prot.pp.clr* with *axis* set to 1; *muon.prot.pp.dsb*). The dsb algorithm uses empty droplets to identify protein-specific noise originating from unbound antibodies^38^. Empty droplets were defined as events that contained between 2 and 3 log_10_(ADT-counts), or (for 5k_pbmc_3p) between 1.5 and 2.8 log_10_(ADT-counts) in the provided unfiltered (“raw”) matrix and did not appear in the filtered matrix. Isotype controls (IgG1, IgG2a and IgG2b) were then used to remove cell-specific noise, as described^38^. Methods were evaluated using UMAPs and Leiden clustering computed as described above, using normalized expression of the 13 proteins profiled in all batches (CD3, CD4, CD8a, CD14, CD16, CD19, CD25, CD45RA, CD45RO, CD56, CD127, PD-1, and TIGIT). Nearest-neighbors were computed using “spearmanr” as the metric, as ADT expression was not scaled (to best-preserve each integration method). Batch integration was quantified with adjusted rand index (ARI) (*sklearn.metrics.adjusted_rand_score*). To test whether landmark registration was robust to changing cell proportions, landmark registration was also performed after removing 95% of the events with positive expression of CD14 (defined as above 2 log(CP1k+1)). The effect of landmark registration on MMoCHi classification was tested by applying MMoCHi to classify subsets across batches to log(CP1k+1)-transformed or landmark registered protein expression, with thresholds manually defined for all batches together or each batch separately (Supplementary Table 7).

### Analysis of organ donor sequencing

CITE-seq data obtained from tissue immune cells was filtered separately. Although hashtag demultiplexing removes inter-sample multiplets, this method cannot detect multiplets between cells from the same tissue site. Thus, multiplets were detected by Scrublet^74^ (*scrublet.Scrublet.scrub_doublets*), using default settings, an expected doublet rate of 0.015, and applying separately to each library-tissue combination of over 100 events. Cells were then filtered to remove events with fewer than 1000 unique counts, fewer than 600 genes detected, or over 10% hemoglobin counts. We used a cluster-based method to remove events either identified as doublets by Scrublet or with high mitochondrial counts, similar to the previously described percolation method^17^. Briefly, a PCA on highly variable genes was calculated, integrated using Harmony^75^ (*scanpy.external.pp.harmony_integrate*), and used for nearest neighbor calculation and Leiden clustering with a resolution of 20. Clusters with significantly higher Scrublet scores (above 0.1) or percent mitochondrial counts (above 15%) were removed. Individual events not captured by these clusters that were identified as Scrublet doublets or had over 25% mitochondrial counts were also removed.

We devised a MMoCHi hierarchy (Fig. 3d; Supplementary Table 8) and performed classification using the algorithm above. Due to insufficient numbers, T_SCM_ were not included in the hierarchy (Supplementary Table 9). Classification fit across a varying number of random forest estimators (trees) was evaluated for prediction accuracy. Overall prediction accuracy was measured using internally held-out events (events that were high-confidence thresholded but excluded from the training of random forest classifiers). Prediction accuracy was also calculated for individual classification nodes using the weighted average of precision, recall, and F1 scores for each child subset using internally held-out, high-confidence events. Gini impurity-based feature importances were automatically calculated during random forest training by scikit-learn. Manual annotation and UMAP calculation were performed on the totalVI latent space, as described above. The clustered dendrogram of classified subsets in each donor was calculated (*scanpy.tl.dendrogram*) on the totalVI latent space with optimal ordering enabled.

Pre-trained classifiers were applied to organ donor PBMC events that were classified as one of: CD4^+^ Naive, CD4^+^ T_CM_, CD4^+^ T_EM_, CD8^+^ Naive, CD8^+^ T_CM_, CD8^+^ T_EM_, CD8^+^ T_EMRA_, or classical monocytes. MMoCHi or Garnett models trained on the sorted FACS dataset (FACS-trained MMoCHi, FACS-trained MMoCHi GEX, and FACS-trained Garnett) or pre-trained ImmClassifier or CellTypist models were applied as described above. For ImmClassifier, CellTypist, and Garnett, Alternative, Unknown, or Erroneous classifications were quantified as described above.

To display of features associated with each subset (Extended Data Fig. 9), the top important features with a log_2_(fold-enrichment) > 2 for that subset. For identification of transcriptomic markers for Naive T cells and T_CM_ (Fig. 5), MMoCHi models were retrained at the CD4^+^ and CD8^+^ Naive/T_CM_ level, given the same multimodal high-confidence thresholding, but trained on only the GEX matrix. To ensure these GEX markers were not learned through tissue-specific contamination, additional models were trained using only T cells from organ-donor blood, and features displayed had to be in the top 1% of important features in both models. To improve robustness, they were additionally filtered to only those with a greater than 10% change in dropout rate.

### Application of MMoCHi to other modalities

To apply MMoCHi to other modalities beyond CITE-seq, we used three public datasets: Ab-seq of sorted T and NK cells^49^, scRNA-seq of a glioma biopsy^50^, and Xenium of a human lymph node obtained from 10x Genomics. For the Ab-seq dataset, only resting cells were used. The gene expression matrix was normalized to log(counts*1,000/total_counts+1), and the protein expression matrix was normalized to log(counts*100/total_counts+1). Clustering was performed on a totalVI latent space, calculated as described above. Manual annotations of these clusters were highly congruent with published annotations.^49^ In the glioma dataset, the gene expression matrix was normalized to log(counts*10,000/total_counts+1). The ratio of the average expression of genes on chromosome 7 to chromosome 10 was used as an additional marker, effectively segregating malignant cells, as described^50^. Manual annotations and UMAP coordinates were obtained from the published clustering analysis. In the Xenium dataset, the gene expression matrix was normalized to log(counts*100/total_counts+1). Gene-expression probes marked as “negative control” or “blank” were removed. The dataset was filtered to remove any events with fewer than 10 total counts and 5 unique genes expressed. Morphological features were calculated using the nucleus and cell boundaries, as determined by the Xenium Onboard Analysis pipeline v1.5.0. Nuclear or cellular area was calculated in μm^2^ by converting boundary vertices to Polygon objects using the Shapely package v2.0.2 (*shapely.geometry.Polygon; shapely.area*). Nuclear or cellular roundness, or circularity, was calculated as the ratio of a shape’s area to the area of the minimum bounding circle of that shape (*shapely.minimum_bounding_circle*). Nuclear DAPI intensity was calculated by obtaining average pixel-intensity of the provided 2D maximum intensity projection (“MIP”) image. To obtain the pixels overlapping with the nucleus, we used the Rasterio package v1.3.9 (*rasterio.features.rasterize*) using a transformation (*affine.Affine.scale*) of 0.2125μm per pixel, as defined in image metadata. These features were supplied to MMoCHi without additional scaling. Manual annotation of clusters was performed on the provided graph-based clusters and displayed on the provided graph-based UMAP projection. MMoCHi was applied to all three datasets using hierarchies of known marker genes, proteins, or chromosome-level expression patterns (Supplementary Table 16). For figure generation, Xenium cell boundaries were plotted on top of the provided 2D autofocus projection image of DAPI staining using the same scaling as above. Cell boundaries were shrunk slightly for display to reduce drawing overlap (using the *shapely.geometry.Polygon.buffer* method, with *join_style* “mitre”). Feature importances were obtained as described above.

### Estimation of computational performance

We measured the computational resources required to train and apply MMoCHi classifiers using a predefined hierarchy and thresholds (Fig. 3c; Supplementary Table 8). Comparisons were performed with multiprocessing enabled for random forest training, using 3^rd^ generation Intel Xeon Scalable processors (3.5 GHz) with 32 vCPUs and 32 GiB of RAM (AWS/EC2 c6i.8xlarge instance). Tests at varying event counts were performed by randomly subsampling the dataset prior to classification.

### Statistical analysis

Statistical comparison of classification performance was calculated using a Friedman rank-sum test (*stats::friedman.test* using the stats library v4.3.1), matched by cell-subset, on F1 scores, followed by multiple comparisons using a paired two-sided Wilcoxon signed-rank test (*stats::pairwise.wilcox.test*) with FDR correction. Significance groups for each method are displayed by lettering, where methods not sharing a letter are significantly different (p < 0.05).

Prior to differential expression, the two groups were subsampled to the same number of events and their expression matrices were downsampled to equal total counts. Differential expression was performed (*scanpy.tl.rank_genes_groups*) on log-normalized expression, with Wilcoxon mode, tie correction, and FDR correction enabled. Genes or proteins with |log_2_(fold change)| > 2 and p < 0.05 were identified as significantly differentially expressed.

## DATA AVAILABILITY

CITE-seq data generated in this study are available in NCBI GEO with the accession code GSE229791. Public datasets used for analyses were downloaded online and are available from: 10x Genomics (https://www.10xgenomics.com/resources/datasets), Open Science Framework (https://doi.org/10.17605/OSF.IO/EDCTN), or NCBI GEO with the accession code GSE116621.

## CODE AVAILABILITY

Code for MMoCHi is available at: https://github.com/donnafarberlab/MMoCHi

Code for demultiplexing CITE-seq and hashtag barcodes is available at: https://github.com/simslab/DropSeqPipeline8

---

**Extended Data Figure 1.**
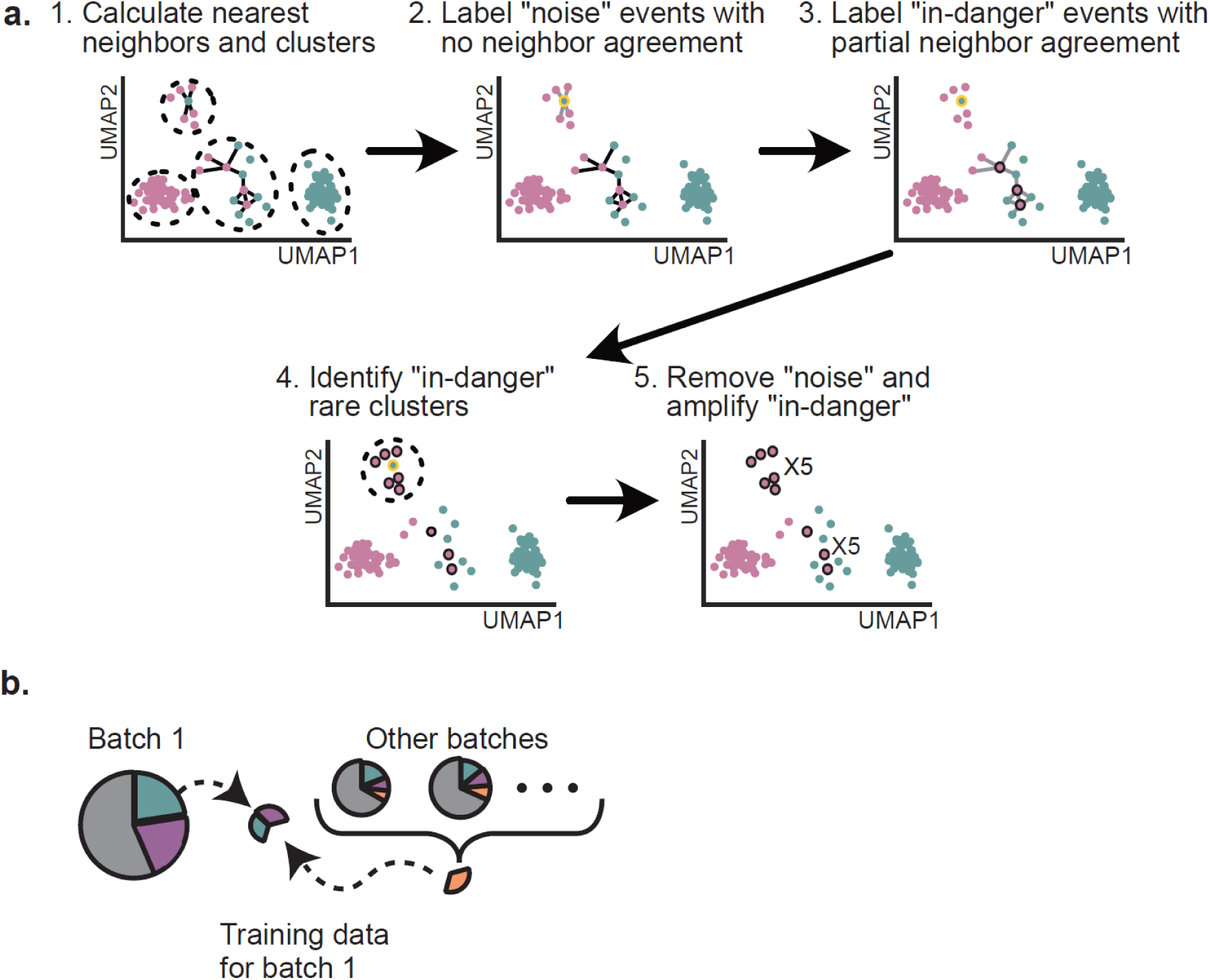
Selection and cleanup of training data. a. Prior to training, high-confidence events were resampled to remove potentially mislabeled events (“noise”) and amplify potentially challenging to classify events (“in danger”). Nearest neighbors are calculated on high-confidence events (1). Events with no neighbors in agreement with their high confidence label are identified as “noise” (2). Events with only some neighbors in agreement or in poorly represented clusters are identified as “in danger” (3,4). Events labeled “noise” are removed from the training data. Events labeled “in danger” are duplicated 5 times in the training dataset (5). b. Training data is selected from a random sample of cleaned-up high-confidence events with rare subsets oversampled so all subsets are equal in the training data. In the case of insufficient training events in individual batches, high-confidence events are spiked into the training data from other batches.

**Extended Data Figure 2.**
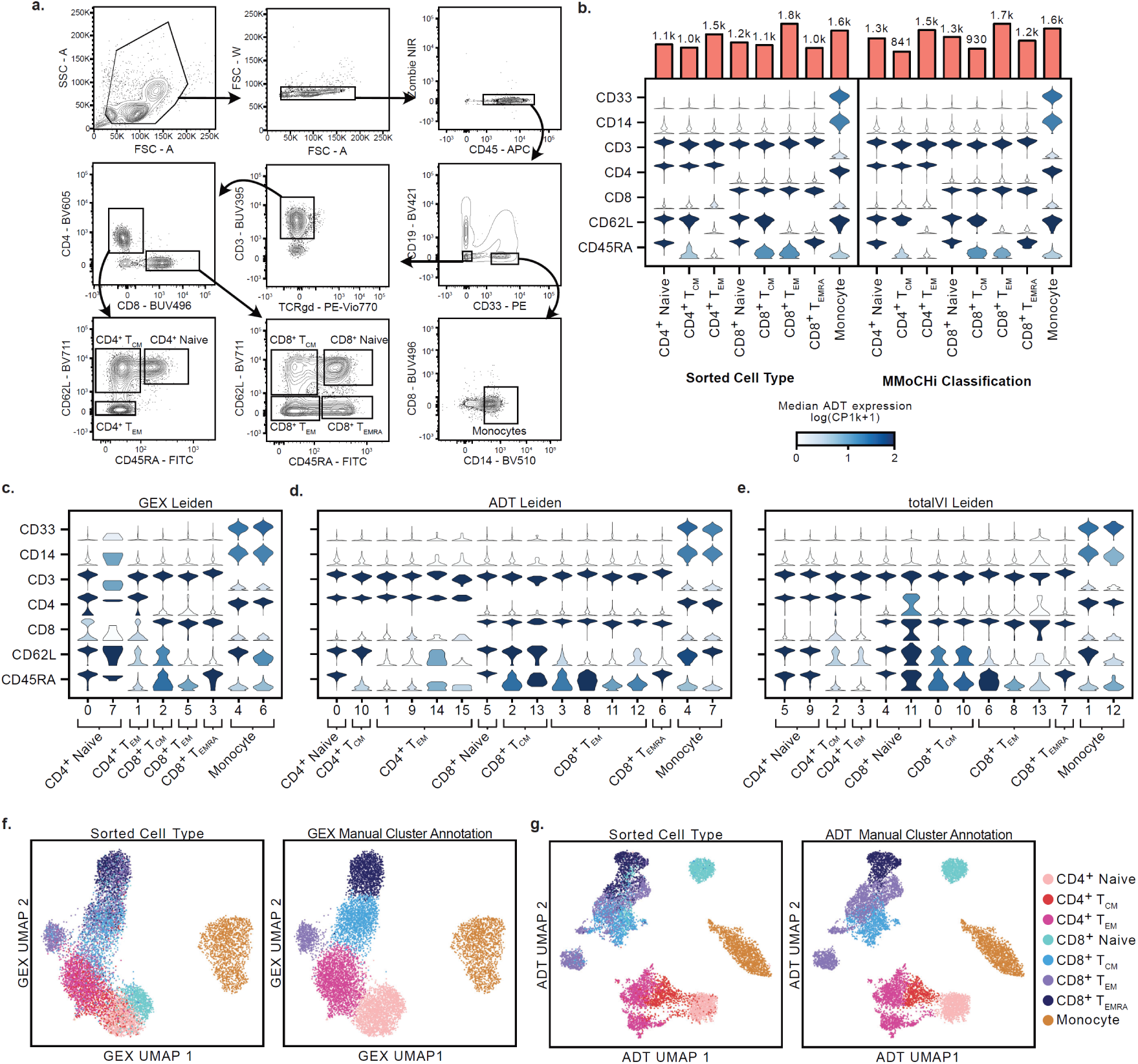
Generation and analysis of MMoCHi validation dataset. a. Gating strategy for fluorescence-activated cell sorting (FACS) of T cell memory subsets and monocytes. Of CD45^+^ cells 6.58% were CD4^+^ Naive, 14.1% were CD4^+^ T_CM_, 5.33% were CD4^+^ T_EM_, 7.01% were CD8^+^ Naive, 3.45% were CD8^+^ T_CM_, 5.14% were CD8^+^ T_EM_, 4.52% were CD8^+^ T_EMRA_, and 17.8% were monocytes. b. Violin plots displaying the distribution of antibody derived tag (ADT) expression for select markers for each sorted population, and each MMoCHi classification. Violins are colored by the median log-normalized ADT counts per thousand. The number of events in each grouping are displayed above the violin plots. c-e. Violin plots displaying distribution of ADT expression for select markers for Leiden clusters of gene expression (GEX; c), ADT expression (d), or totalVI latent space (e). Clusters are grouped by their manual annotation. f-g. UMAPs of principal component analysis used for GEX annotation (f), or ADT annotation (g), colored by sorted cell type (identified by hashtag oligo; HTO) or manual annotation. T_CM_, central memory T cell; T_EM_, effector memory T cell; T_EMRA_, terminally differentiated effector memory T cell.

**Extended Data Figure 3.**
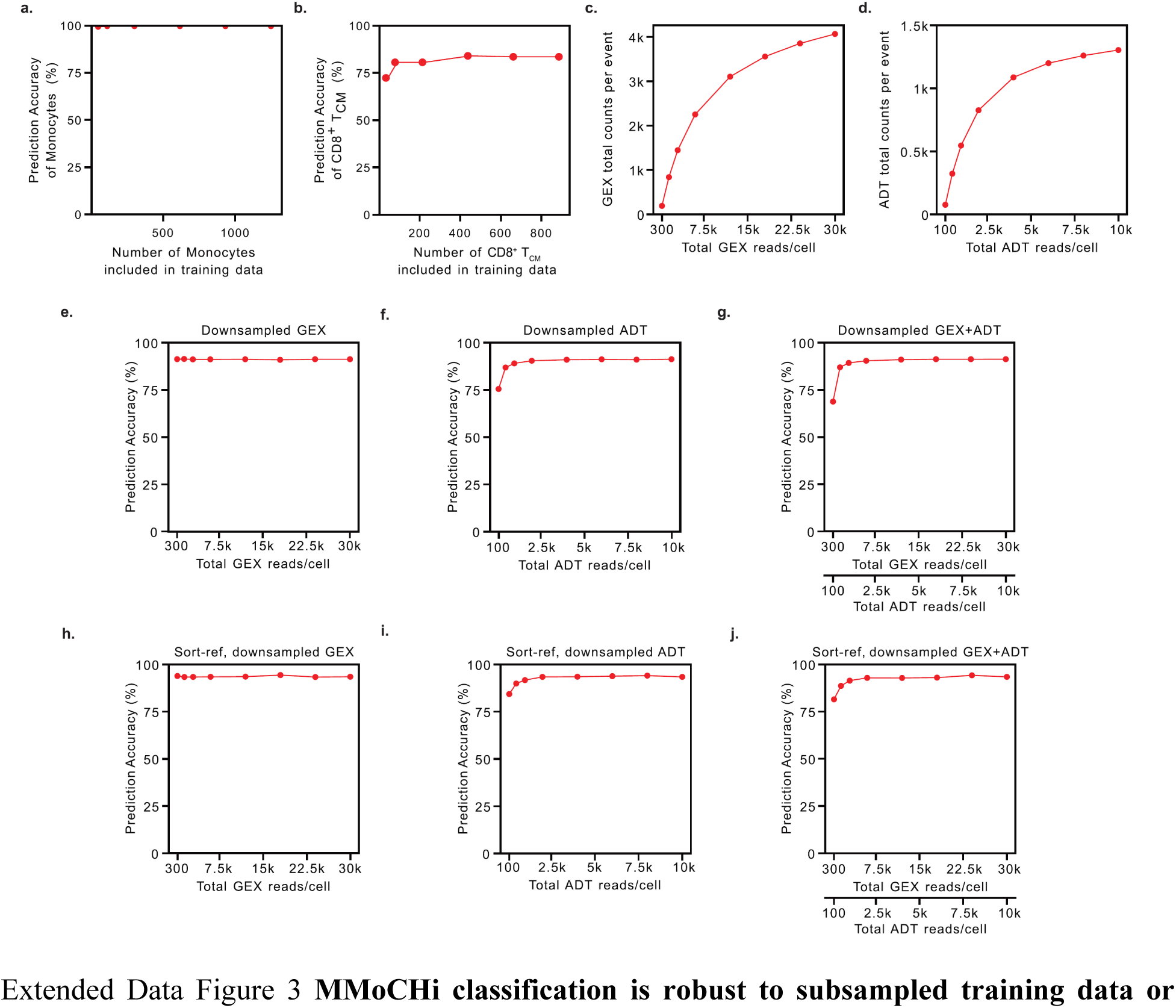
MMoCHi classification is robust to subsampled training data or downsampled reads. a-b. Classification accuracy of a monocytes (a) or CD8^+^ T_CM_ following subsampling of each subset prior to training data selection (only 40-50% of which are identified by high-confidence thresholding). Sorted cell type labels (derived from hashtag-oligo; HTO) were used as “truth” and accuracy was calculated using a held-out data. c-d. Line plots depicting the effect of downsampling aligned reads of gene expression (GEX; c) or antibody derived tag (ADT) expression (d) on the average total counts in the expression matrix per event. c-h. Balanced classification accuracy of sorted T cell memory subsets and monocytes, with MMoCHi classifiers trained using high-confidence thresholding on downsampled events (e-g) or trained using the HTO-derived sort labels as reference and tested on held-out data (h-j). Classifiers were exposed to downsampled GEX and normal ADT expression (e,h), normal GEX and downsampled ADT expression (f,i), or downsampled GEX and ADT expression (g,j) for both training and prediction.

**Extended Data Figure 4.**
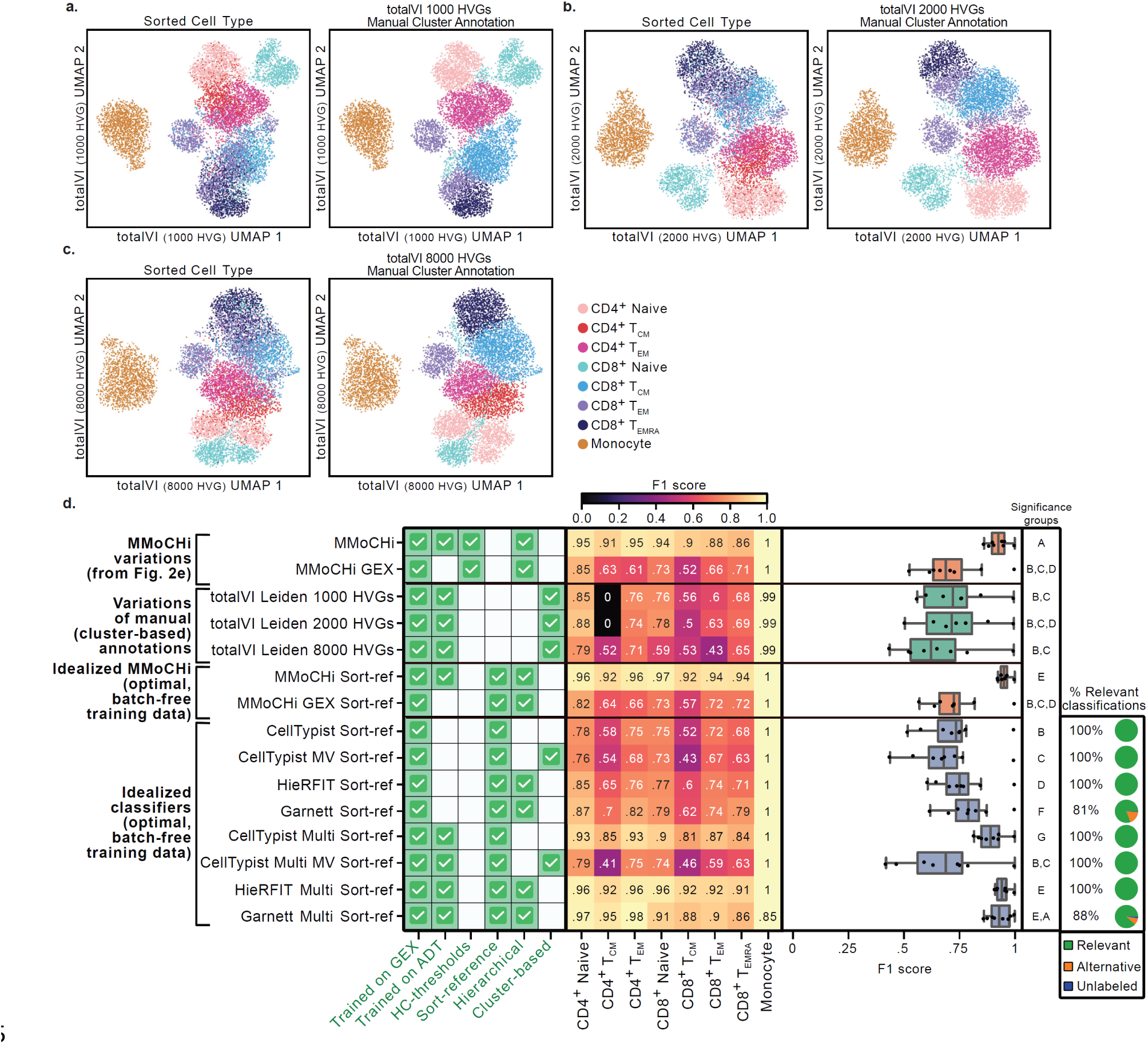
MMoCHi benchmarked against other variations and controls. a-c. UMAPs of totalVI embeddings calculated using various numbers of most highly variable genes colored by sorted cell type (left) or manual cluster-based annotation (right). d. Performance comparison using F1 scores, calculated for each cell subset using HTO-derived sorted cell type labels as truth. F1 scores for each method were aggregated in box and whisker plots (right). Features of each method are labeled for context (left) using green checks: “Trained on GEX/ADT”—whether gene expression (GEX) or antibody derived tag (ADT) expression data were used for model training. “HC-thresholds”—whether high-confidence thresholding was used for training data selection. “Sort-reference”—whether HTO-derived sorted cell type labels were used to train the classifier. “Hierarchical”—whether annotation was performed on multiple levels. “Cluster-based”—whether annotations were applied to unsupervised clusters. For scRNA-seq classifiers, the percent of classifications that were relevant to this analysis, were alternative (but potentially valid) classifications or were unlabeled are shown using pie charts (far right). Statistical significance was calculated using a Friedman rank-sum test, matched by subset, followed by multiple comparisons using paired Wilcoxon signed-rank tests and FDR correction. Significance groups for each method are shown by lettering, where methods not sharing a letter are significantly different (p < 0.05).

**Extended Data Figure 5.**
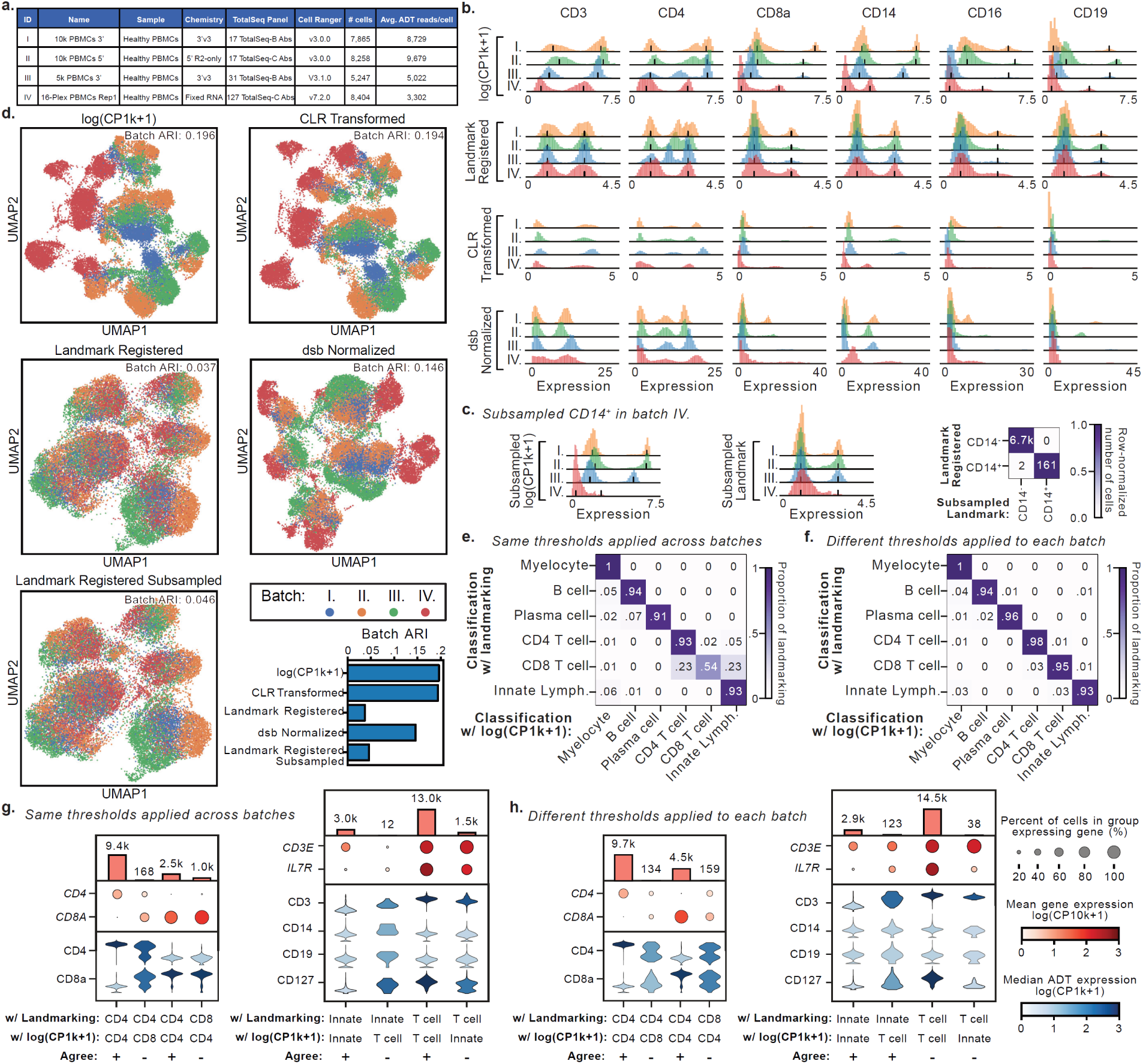
Landmark registration integrates ADT expression and allows for consistent thresholding. a. Metadata for each dataset highlighting features that may contribute to batch effects. b. Histograms displaying normalized Antibody Derived Tag (ADT) expression distribution for selected proteins. Histograms for each batch are grouped by normalization technique. Peaks identified for landmark registration are displayed on the log(CP1k+1)-transformed expression. Aligned peaks are displayed on the landmark registered expression. c. Histograms as in (b) with the number of CD14^+^ cells in batch IV subsampled to 5%. Heatmap displaying the number of CD14^+^ cells by landmark-registration before and after subsampling. d. UMAPs of the expression of the 16 ADTs in all panels, colored by batch to display integration. Adjusted Rand Index (ARI) measures similarity of clustering and batch identity, where lower values indicate stronger integration. e-f. Row-normalized heatmaps displaying MMoCHi classifications (see Supplementary Table 7 for hierarchy specification) when run using landmark-registered or log(CP1k+1)-transformed ADT expression. g-h. Plots depicting expression of selected markers on populations classified by MMoCHi with or without landmark registration. Dot plots display gene expression. Dot size represents the percent of cells in the group expressing a gene, and dots are colored by the mean log-normalized GEX counts per ten thousand. Violin plots display the distribution of antibody derived tag (ADT) expression for each population. Violins are colored by the median log-normalized ADT counts per thousand. The number of cells in each group are displayed above the dot plots. Populations are denoted by a “+” where annotations agree, and a “-” where they disagree. MMoCHi was run twice, with either the same high-confidence thresholds defined across all batches (e,g), or separately for each batch (f,h). CLR: Centered Log-Ratio; dsb: denoised and scaled by background.

**Extended Data Figure 6.**
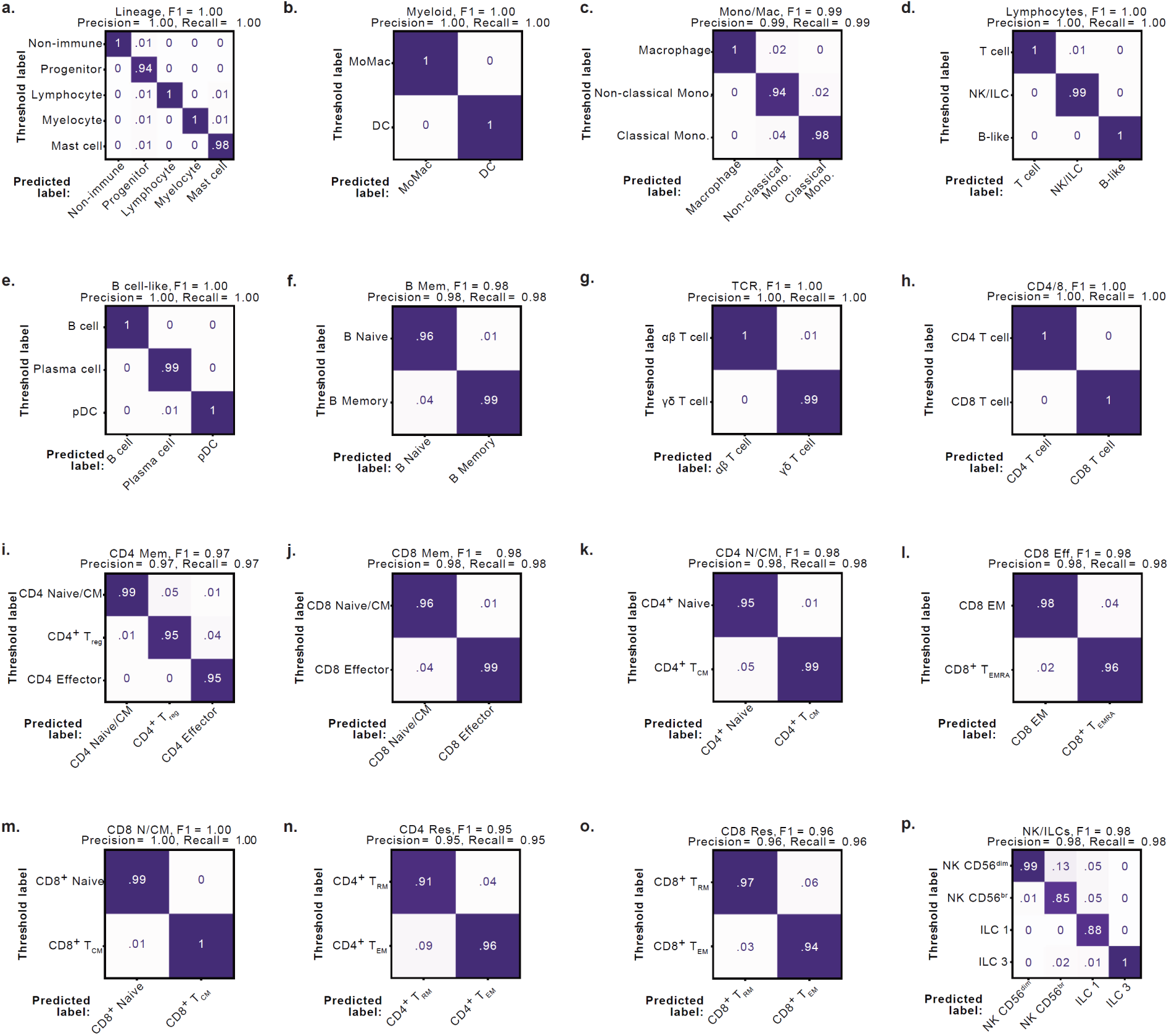
Random forests perform well at each level of MMoCHi hierarchy. a-p. Column-normalized confusion matrices depicting the fit of each random forest classifier in the organ-donor hierarchy (Fig. 3c; Supplementary Table 8). MMoCHi classifications are compared with internally held-out high-confidence threshold labels at each node. T_CM_, central memory T cell; T_reg_, regulatory T cell; T_EM_, effector memory T cell; T_RM_, resident memory T cell; T_EMRA_, terminally differentiated effector memory T cell; ILC, innate lymphoid cell; NK, natural killer cell; pDC, plasmacytoid dendritic cell; DC, dendritic cell; Mono, monocyte.

**Extended Data Figure 7.**
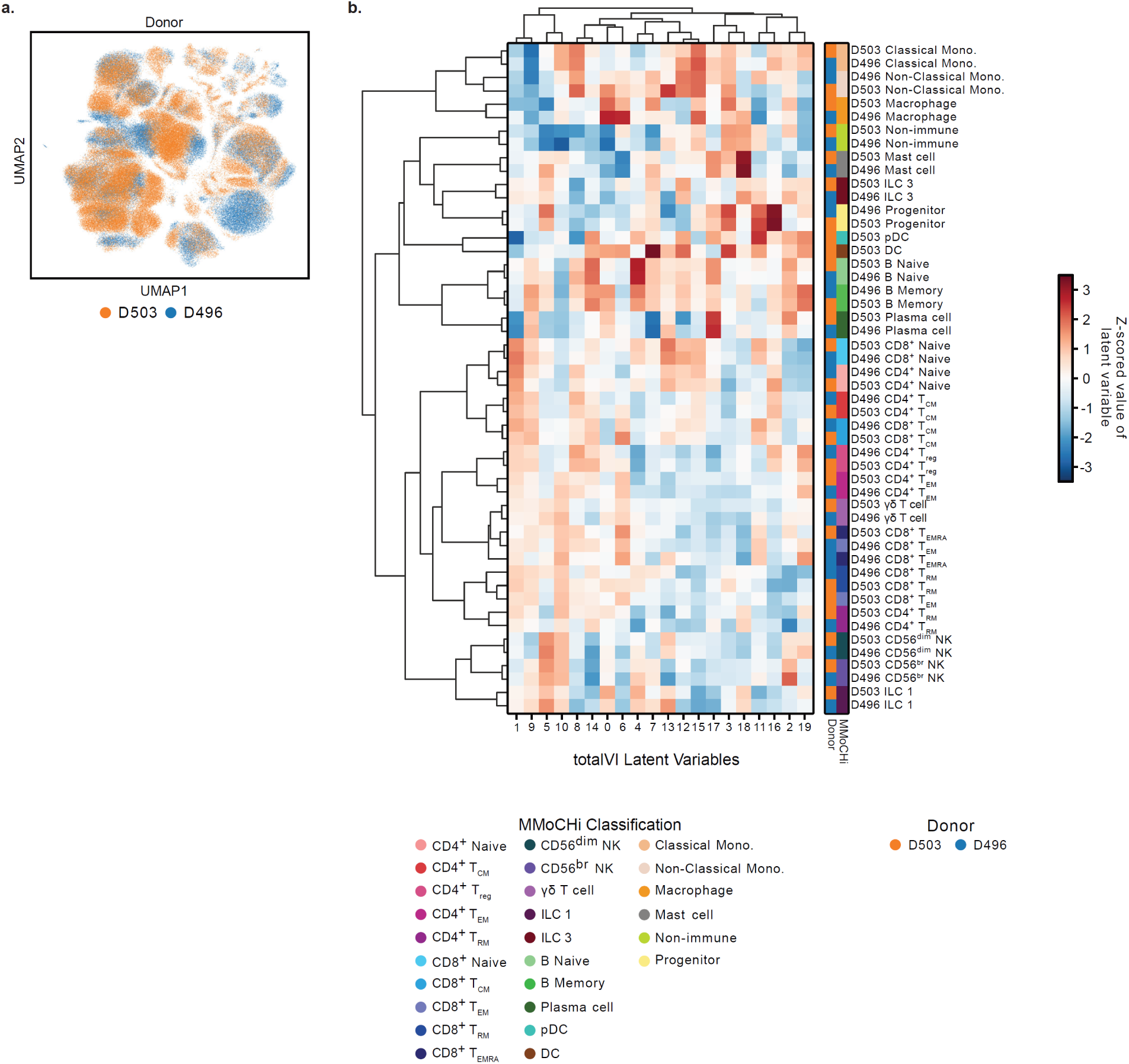
MMoCHi classifies cell types across donor. a. UMAP of donor-integrated totalVI latent space, colored by donor. b. Z-scored expression of totalVI latent variables averaged across each classified cell type in each donor (filtered by donor-subset pairs with at least 100 events). Dendrogram clustering of cell type-donors (left) and factors (top) calculated using Pearson correlation on the totalVI latent space. Rows are labeled with color by donor (left) and MMoCHi classification (right). LNG, lung; BAL, bronchoalveolar lavage; LLN, lung lymph node; SPL, spleen; JEL, jejunum epithelial layer; JLP, jejunum lamina propria; BOM, bone marrow; BLD, blood; T_CM_, central memory T cell; T_reg_, regulatory T cell; T_EM_, effector memory T cell; T_RM_, resident memory T cell; T_EMRA_, terminally differentiated effector memory T cell, NK, natural killer cell; ILC, innate lymphoid cell; DC, dendritic cell; pDC, plasmacytoid dendritic cell; Mono, monocyte.

**Extended Data Figure 8.**
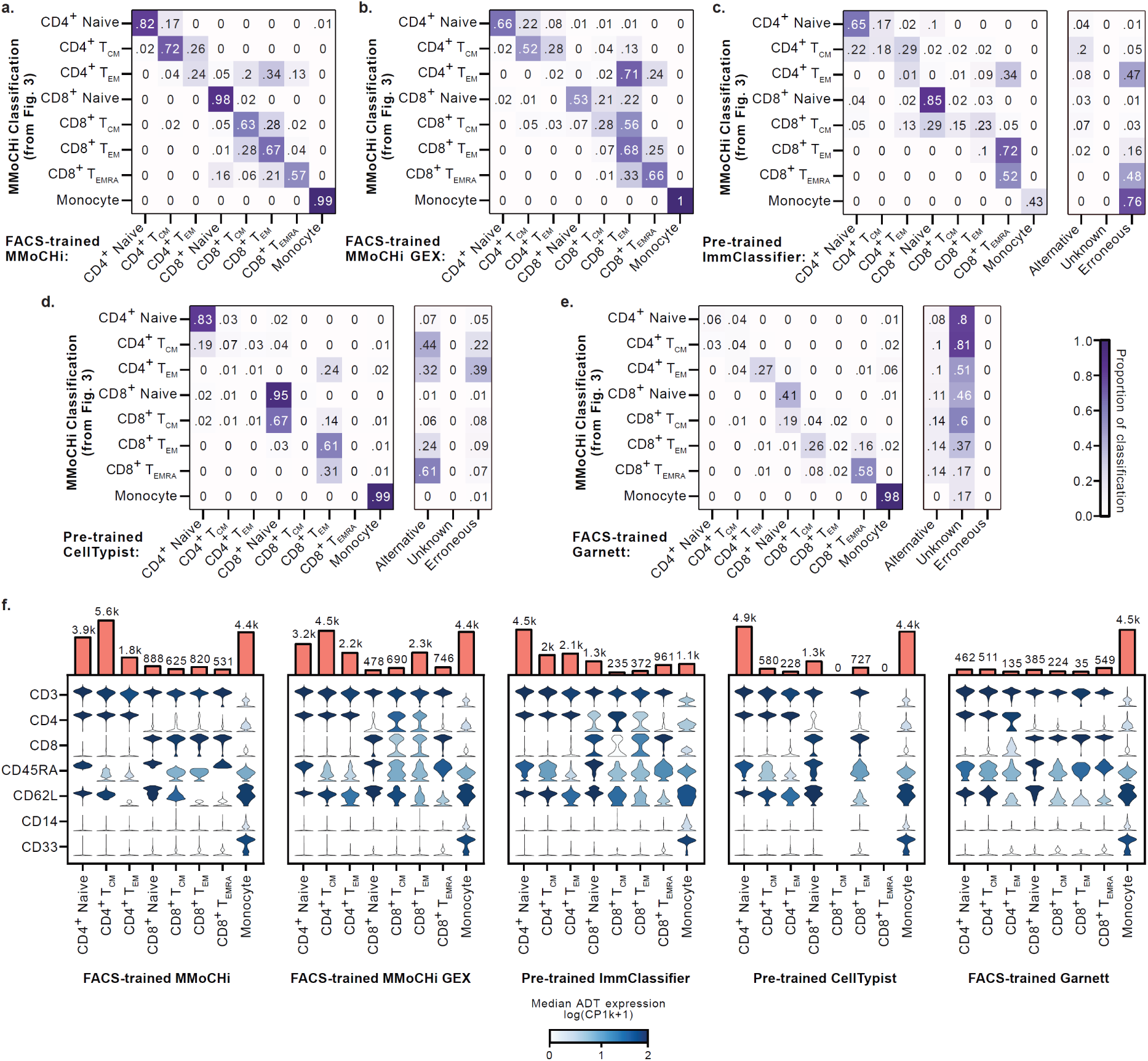
Pre-trained MMoCHi classifiers effectively annotate new data. a–e. Row-normalized heatmaps displaying agreement between pre-trained classifiers and the MMoCHi classification obtained in Fig. 3, subset to only T cell memory subsets and monocytes from blood. Classifiers applied include: MMoCHi trained on sorted blood (FACS-trained; a), MMoCHi trained on sorted blood only using gene expression (FACS-trained GEX; b), ImmClassifier using the provided pre-trained model (c), Celltypist using a provided pre-trained model (d), and Garnett trained on sorted blood (e). Alternative (but potentially valid), Unknown (unlabeled), and Erroneous (cell types not present in sampled dataset) are indicated separately. f. Plots depicting expression of selected cell type markers, on cells grouped by their classification. Violin plots display the distribution of antibody derived tag (ADT) expression for each population. Violins are colored by the median log-normalized ADT counts per thousand. The number of events in each grouping are displayed above the violin plots. T_CM_, central memory T cell; T_EM_, effector memory T cell; T_EMRA_, terminally differentiated effector memory T cell.

**Extended Data Figure 9.**
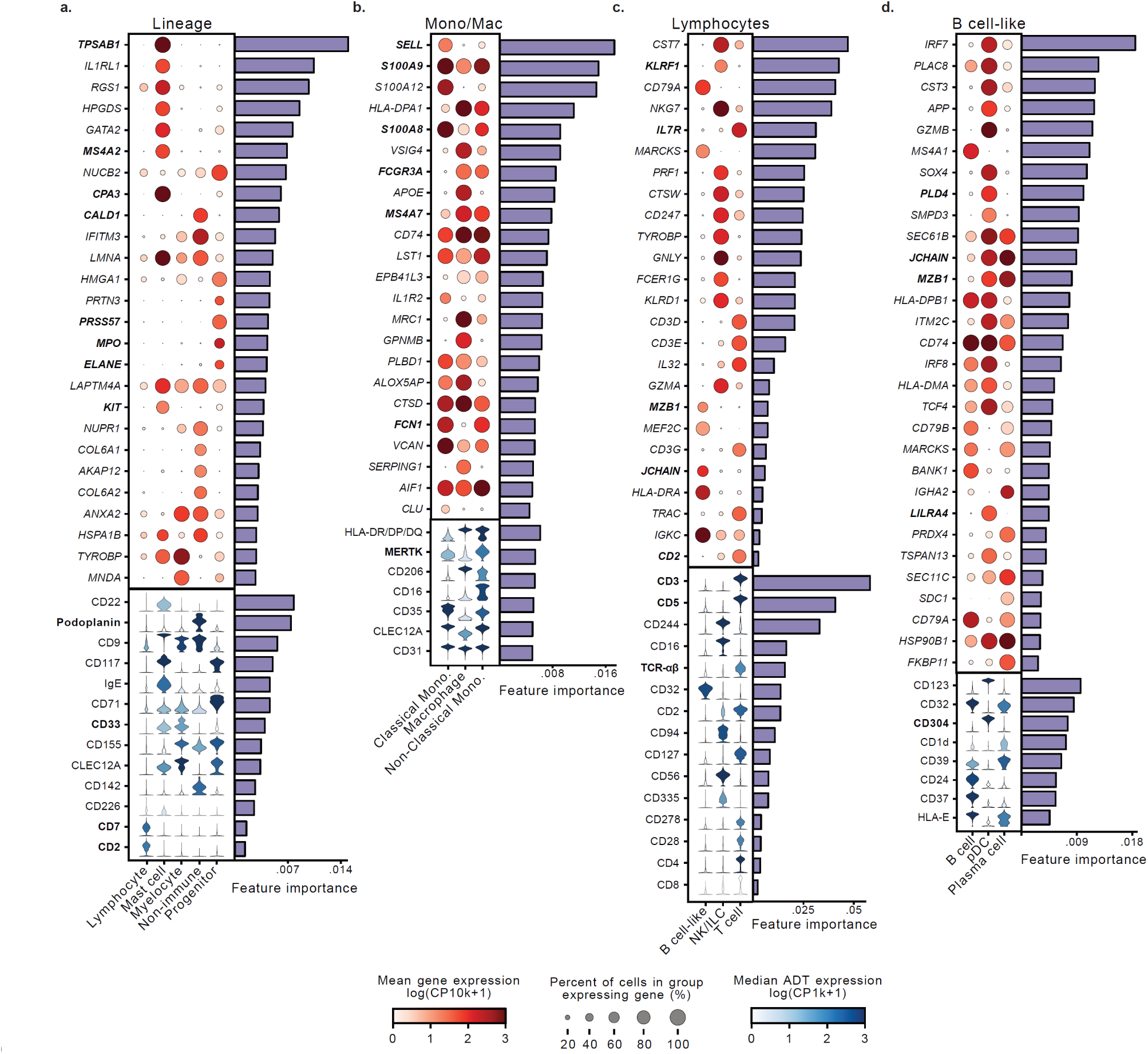
Interpretation of MMoCHi random forests using feature importances reveal immune cell lineage markers. a-d. Expression of important features for classification for the Lineage (a), Mono/Mac (b), Lymphocytes (c), and B cell-like (d) nodes in the organ donor hierarchy (Fig. 3c). The top important features associated with each subset (log_2_(fold change) > 1.5) are displayed to include representation of features specific to each subset. All features displayed are within the top 1% of important features. Features used for high-confidence thresholding are highlighted in bold. Dot plots display gene expression (GEX), where dot size represents the percent of cells in the group expressing a gene, and dots are colored by the mean log-normalized GEX counts per ten thousand. Violin plots display the distribution of antibody derived tag (ADT) expression and are colored by the median log-normalized ADT counts per thousand. The impurity-based importance of each feature displayed is shown in a bar chart to the right. All features displayed were also significantly differentially expressed (p < 0.05). Statistical significance was calculated using a two-sided Wilcoxon with tie correction, followed by a Benjamini–Hochberg adjustment for multiple comparisons. Mac, Macrophage; Mono, monocyte; NK, natural killer cell; ILC, innate lymphoid cell; pDC, plasmacytoid dendritic cell.

**Extended Data Figure 10.**
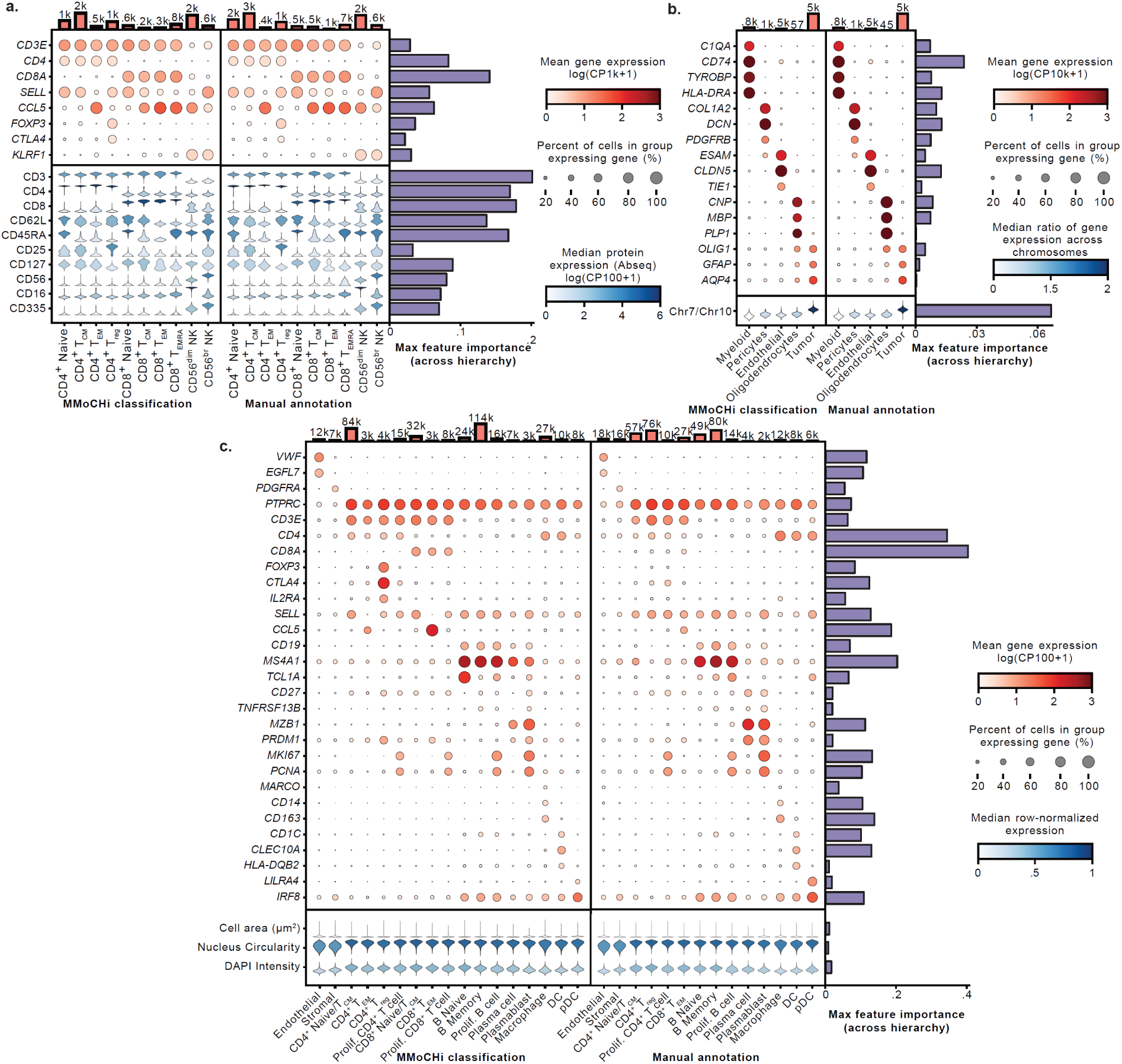
Marker gene and protein expression on manually annotated and MMoCHi classified populations. a-c. Expression of select cell type markers on cells grouped by either their MMoCHi classification (left) or their manual cluster annotation (right) for the Ab-seq (a), Glioma (b), and Xenium (c) datasets. Dot plots display gene expression (GEX). Dot size represents the percent of cells in the group expressing a gene, and dots are colored by the mean log-normalized GEX. Violin plots display the distribution of protein expression (a), the ratio of gene expression across chromosomes (b), or row-normalized physical attributes of cells (c), for each MMoCHi-classified or manually-annotated population. Violins are colored by the median value. T_CM_, central memory T cell; T_reg_, regulatory T cell; T_EM_, effector memory T cell; T_EMRA_, terminally differentiated effector memory T cell, NK, natural killer cell; DC, dendritic cell; pDC, plasmacytoid dendritic cell; Prolif., Proliferating.

