## Supplementary Figures and Table Legends for "Multimodal hierarchical classification of CITE-seq data delineates immune cell states across lineages and tissues"

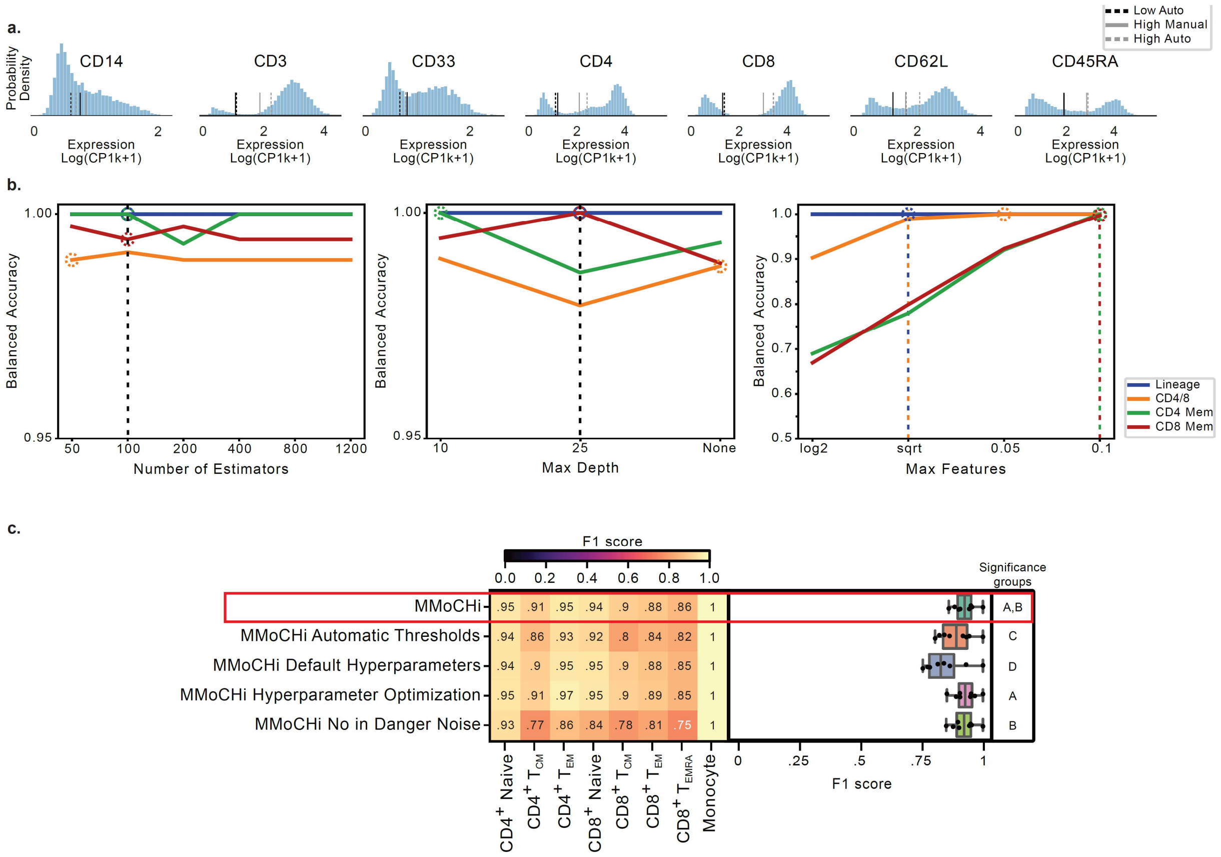


Supplementary Figure 1 **MMoCHi is robust to automatic thresholds and hyperparameter optimization.** a. Histogram displaying normalized expression of each protein used in classification of sorted T cell memory subsets and monocytes, with dashed lines indicating the automatic thresholds and solid lines displaying manually selected thresholds. Positive thresholds are displayed in gray, negative thresholds are displayed in black. b. Line plots colored by classification node depicting classification performance using various hyperparameter values compared to high confidence thresholds. Manually selected hyperparameters used for training at each classification node (dashed lines) and automatically selected optimal hyperparameters (dashed circles) are shown. "Max depth"—The maximum depth of trees in random forest; "Max features"—The fraction of total features presented when looking for the best split during training; "Number of Estimators"—The number of decision trees trained at each classification node c. Performance comparison using F1 scores, calculated for each cell subset using HTO-derived sorted cell type labels as truth. F1 scores for each method were aggregated in box and whisker plots. Statistical significance was calculated using a Friedman rank-sum test, matched by subset, followed by multiple comparisons using paired Wilcoxon signed-rank tests and FDR correction. Significance groups for each method are shown by lettering, where methods not sharing a letter are significantly different ($p < 0.05$). log2: $\log_2$(total features), sqrt: √(total features); $T_{CM}$, central memory T cell; $T_{EM}$, effector memory T cell; $T_{RM}$, resident memory T cell; $T_{EMRA}$, terminally differentiated effector memory T cell; Mono, monocyte.

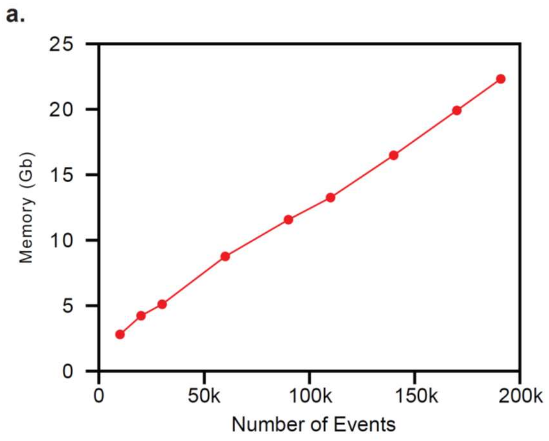


Supplementary Figure 2 **Time and memory performance of MMoCHi classification.** a-b. Memory usage (a) and run time (b) to classify 26 subsets using CITE-seq of two human organ donors across multiple tissues. Tests were performed using a predefined hierarchy and thresholds with multiprocessing enabled for random forest training on a computer with 3$^{rd}$ Generation Intel Xeon Scalable processors (3.5 GHz), with 32 vCPUs and 32 GiB of RAM. Tests at different event counts were performed with random subsampling prior to classification.

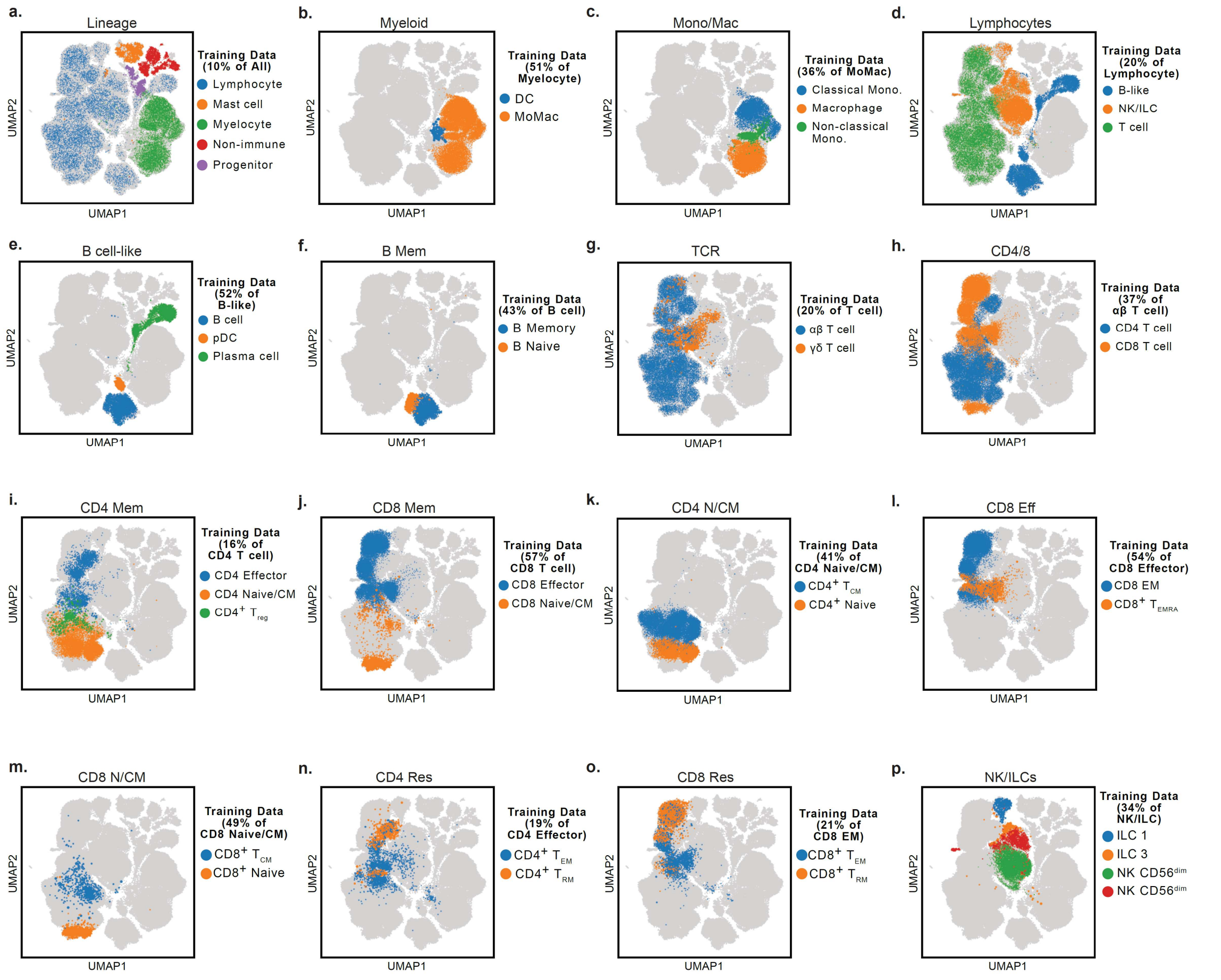


Supplementary Figure 3 **Organ donor training data.** a-p. Training data selected at each classification node of the organ donor hierarchy (Fig. 3d). UMAPs of totalVI latent space, where events used for training are colored by their high-confidence threshold label. Dot size is proportional to the percent of the training dataset they represented after resampling of training data (see Extended Data Fig. 1). The total percent of events represented in the training dataset is listed for each classification node. $T_{CM}$, central memory T cell; $T_{reg}$, regulatory T cell; $T_{EM}$, effector memory T cell; $T_{RM}$, resident memory T cell; $T_{EMRA}$, terminally differentiated effector memory T cell; ILC, innate lymphoid cell; NK, natural killer cell; pDC, plasmacytoid dendritic cell; DC, dendritic cell; Mono, monocyte

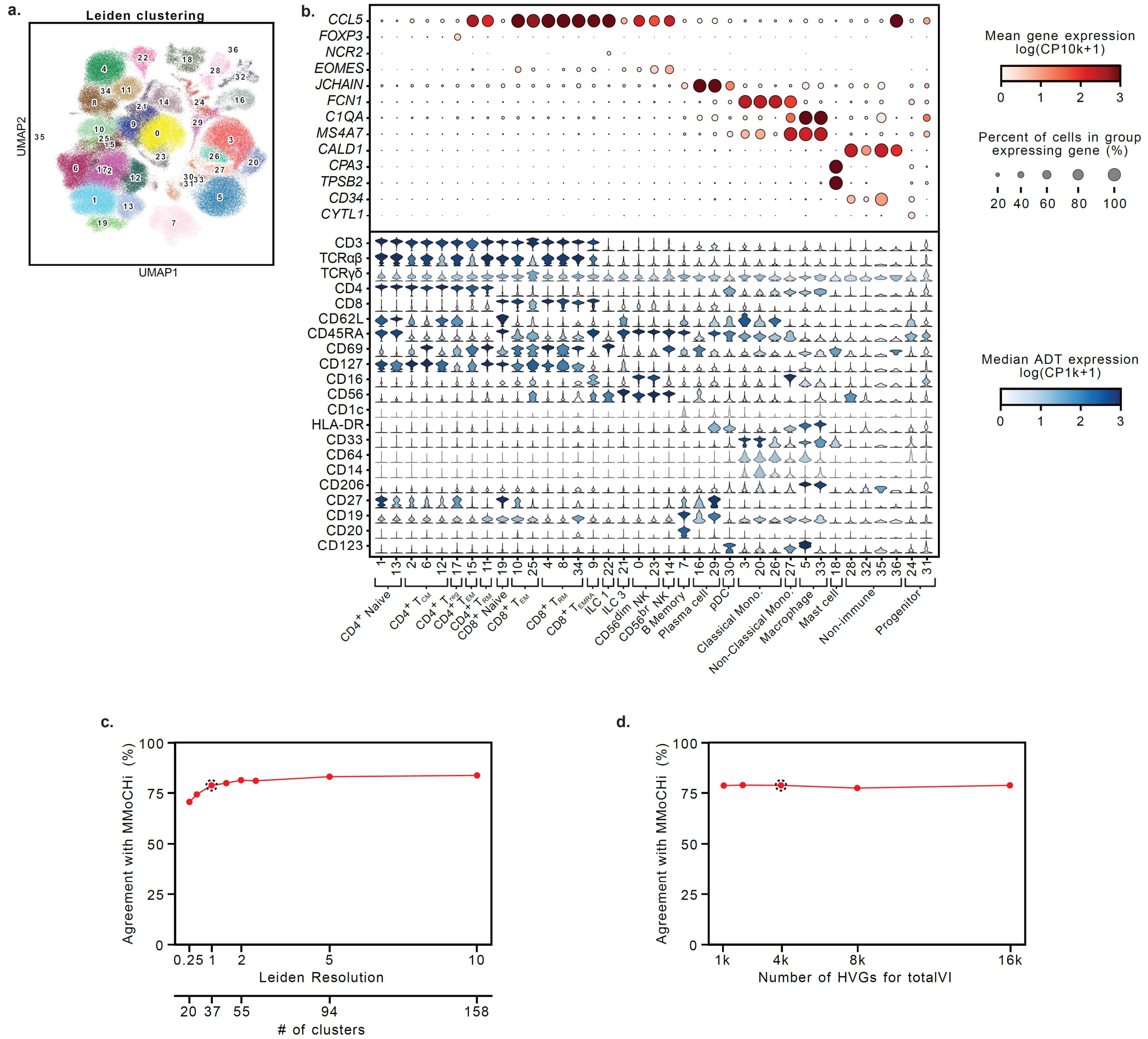


Supplementary Figure 4 **Manual annotation of Leiden clusters by marker gene and protein expression.** a. UMAP of totalVI latent space displaying Leiden clusters. b. Expression of select markers for each cluster, grouped by manually annotated cell type. Dot plots display gene expression (GEX). Dot size represents the percent of cells in the group expressing a gene, and dots are colored by the mean log-normalized GEX counts per ten thousand. Violin plots display the distribution of antibody derived tag (ADT) expression for each cluster. Violins are colored by the median log-normalized ADT counts per thousand. c-d. Line plots depicting agreement between manual annotation and MMoCHi classification when increasing Leiden resolution (c) or variation in number of highly variable genes (HVGs) used to create totalVI latent space (d). Dashed circle indicates Leiden resolution or number of highly variable genes used for analysis (Fig. 3-4).

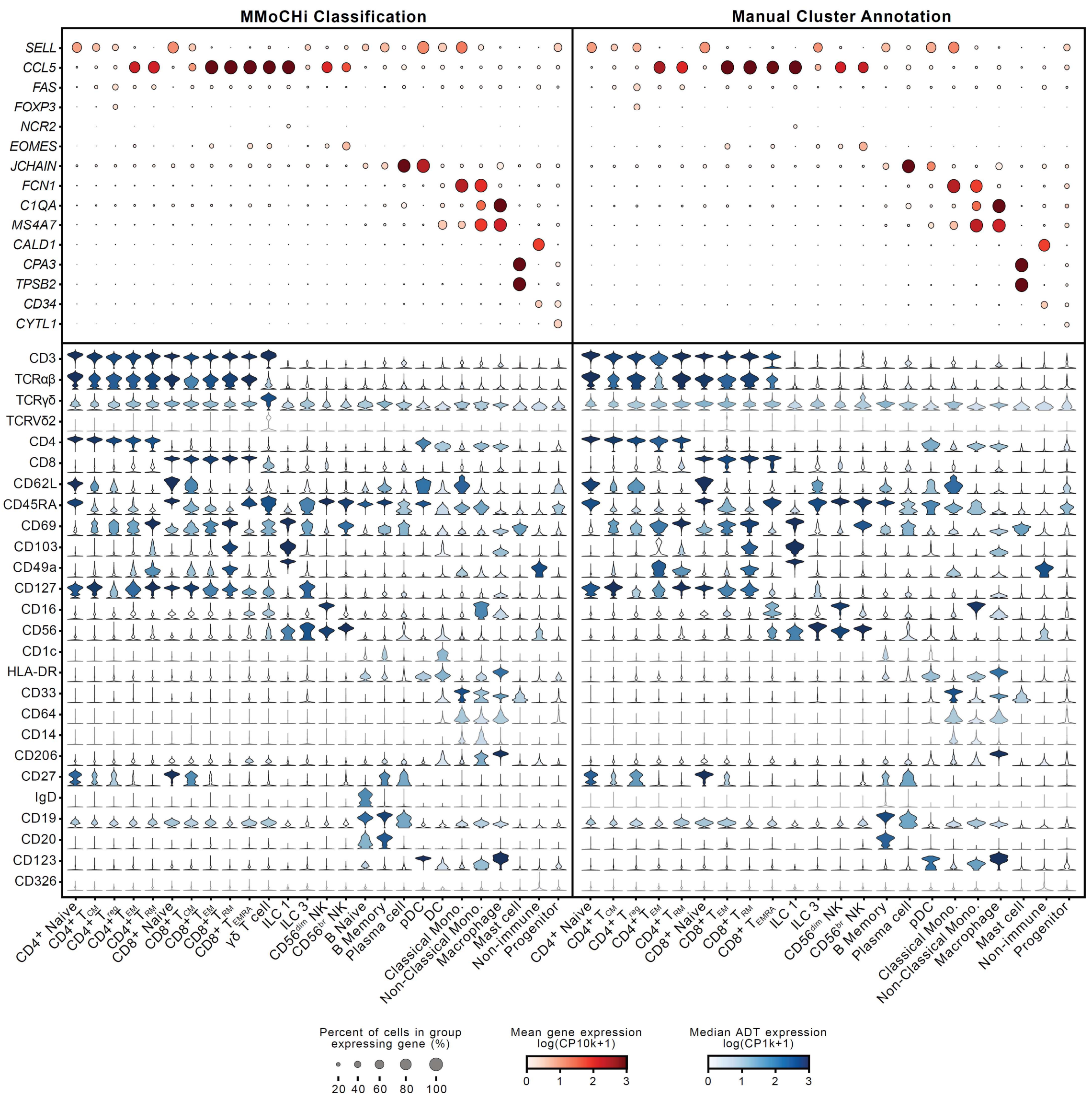


Supplementary Figure 5 **Marker gene and protein expression on manually annotated and MMoCHi classified populations.** Expression of select cell type markers on cells grouped by either their MMoCHi classification (left) or their manual cluster annotation (right). Dot plots displaying gene expression (GEX). Dot size represents the percent of cells in the group expressing a gene, and dots are colored by the mean log-normalized GEX counts per ten thousand. Violin plots display the distribution of antibody derived tag (ADT) expression for each MMoCHi-classified or manually-annotated population. Violins are colored by the median log-normalized ADT counts per thousand. $T_{CM}$, central memory T cell; $T_{reg}$, regulatory T cell; $T_{EM}$, effector memory T cell; $T_{RM}$, resident memory T cell; $T_{EMRA}$, terminally differentiated effector memory T cell; ILC, innate lymphoid cell; NK, natural killer cell; pDC, plasmacytoid dendritic cell; DC, dendritic cell; Mono, monocyte

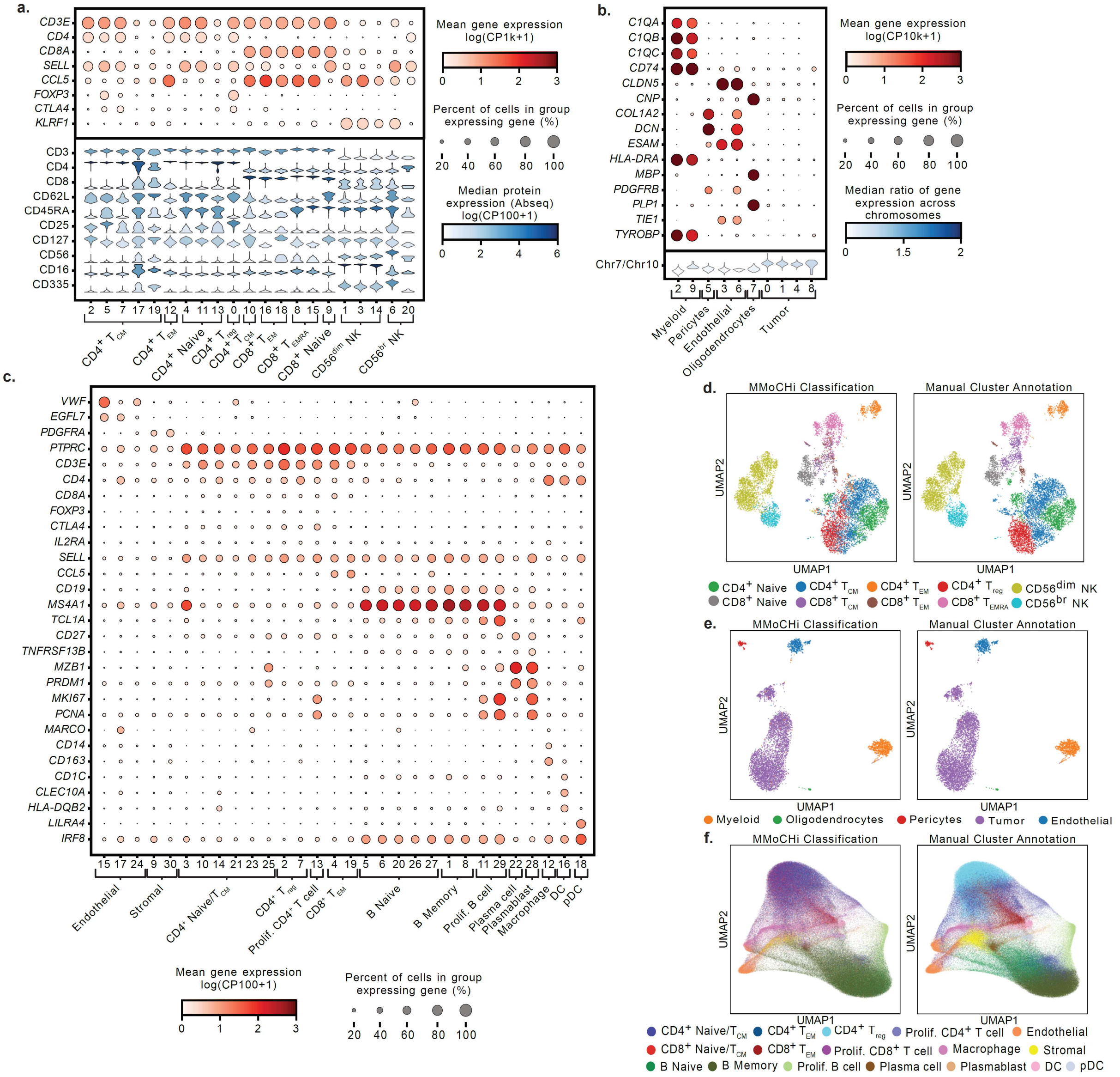


Supplementary Figure 6 **Manual annotation of Leiden clusters by marker expression.** a-c. Expression of select markers for each cluster, grouped by manually annotated cell type for the Ab-seq (a), Glioma (b), and Xenium (c) datasets. Dot plots display gene expression (GEX). Dot size represents the percent of cells in the group expressing a gene, and dots are colored by the mean log-normalized GEX. Violin plots display the distribution of protein expression (a) or the ratio of gene expression across chromosomes (b) for each annotated cluster. Violins are colored by the median expression. d-f. UMAPs of the Ab-seq (d), Glioma (e), and Xenium (f) datasets colored by MMoCHi classification, or manual cluster annotations. $T_{CM}$, central memory T cell; $T_{reg}$, regulatory T cell; $T_{EM}$, effector memory T cell; $T_{EMRA}$, terminally differentiated effector memory T cell, NK, natural killer cell; DC, dendritic cell; pDC, plasmacytoid dendritic cell; Prolif., Proliferating

### SUPPLEMENTARY TABLE LEGENDS

Supplementary Table 1 **Donor metadata** Metadata for the organ donors (D496 and D503) and the blood donor (Blood Donor A) used in this study.

Supplementary Table 2 **Antibodies** A list of antibody clones and sources used in the FACS panel, custom TotalSeq-A CITE-seq panel, and for hashtag-oligo (HTO) staining. Where applicable, the conjugated fluorophore or barcode sequence used for alignment are included.

Supplementary Table 3 **Sort purity** Sort purity of T cell subsets and monocytes after FACS. Purity is represented as the percentage of singlet events falling within the sorted subset's gates using the same gating strategy as the sort (Extended Data Fig. 2a).

Supplementary Table 4 **MMoCHi hierarchy (FACS validation)** Subset definitions and thresholds used to construct the MMoCHi hierarchy. Hierarchical relationships are defined by subset parents (subsets with no parent represent the first classification level). High-confidence events for each subset are selected as events matching the listed marker expression (above the positive threshold for all "Positive" markers, and below the negative threshold for all "Negative" markers). Thresholds are defined for the listed protein markers in log(CP1k + 1).

Supplementary Table 5 **Annotation performance metrics** F1 score, Precision, Recall, and Accuracy of MMoCHi and various annotation methods compared to sorted population labels (Fig. 2), as defined by hashtag expression (see Methods). For tools that produced non-relevant classifications, the total proportion of alternative, unlabeled, or erroneous classifications are quantified (see Methods).

Supplementary Table 6 **Garnett definitions** Garnett marker file providing the hierarchy and gene expression marker-based definitions used for classification. See Methods for details on marker selection.

Supplementary Table 7 **MMoCHi hierarchy (landmark registration)** Subset definitions and thresholds used to construct the MMoCHi hierarchy. Hierarchical relationships are defined by subset parents (subsets with no parent represent the first classification level). High-confidence events for each subset are selected as events matching the listed marker expression (above the positive threshold for all "Positive" markers, and below the negative threshold for all "Negative" markers). All gene expression markers are suffixed with "_gex". Thresholds for protein markers are defined using log(CP1k + 1) or landmark-registered expression, as denoted. Thresholds for gene expression markers in log(CP10k + 1).

Supplementary Table 8 **MMoCHi hierarchy (organ donors)** Subset definitions and thresholds used to construct the MMoCHi hierarchy. Hierarchical relationships are defined by subset parents (subsets with no parent represent the first classification level). High-confidence events for each subset are selected as events matching the listed marker expression (above the positive threshold for all "Positive" markers, and below the negative threshold for all "Negative" markers). All gene expression markers are suffixed with "_gex". Thresholds for protein markers are defined using landmark-registered expression (see Methods). Thresholds for gene expression markers in log(CP10k + 1).

Supplementary Table 9 **Insufficient availability of $T_{SCM}$ in organ donor dataset for classification** Total number of cells across D496 and D503 that could be defined as high-confidence CD4+ or CD8+ T cells by protein gates, and percentage of those cells that could be identified as $T_{SCM}$. Thresholds for each marker are defined in Supplementary Table 7.

Supplementary Table 10 **Cross-dataset annotation performance metrics** F1 score, Precision, Recall, and Accuracy of pre-trained MMoCHi and other pre-trained classifiers on organ donor PBMCs compared to MMoCHi classification labels (Fig. 4), as defined by hashtag expression (see Methods). For tools that produced non-relevant classifications, the total proportion of alternative, unlabeled, or erroneous classifications are quantified (see Methods).

Supplementary Table 11 **Important features at each classification level** Gini impurity-based importance for each feature used for classification at each level of the organ donor classification, used to construct Fig. 5. Feature names for each level are suffixed with "_mod_GEX" for gene expression and "_mod_landmark_protein" for landmark-registered protein expression.

Supplementary Table 12 **Differential expression between classified immune cell subsets** Differential expression of gene expression (GEX) or antibody-derived tag (ADT) expression between each MMoCHi-classified subset listed in Fig. 5 and all other subsets at the same classification level.

Supplementary Table 13 **Important features for Naive/TCM classification** Gini impurity-based importance for each feature used for $CD4^+$ and $CD8^+$ Naive/$T_{CM}$ classification, used to construct Fig. 6c-d. Feature names for each level are suffixed with "_mod_GEX" for gene expression. Classification was performed on T cells from all donor tissue sites (cd4_ncm, cd8_ncm) and on only T cells from organ donor blood (cd4_ncm_bld, cd8_ncm_bld).

Supplementary Table 14 **Differential expression of $CD4^+$ and $CD8^+$ Naive vs $T_{CM}$** Differential expression of gene expression (GEX) between MMoCHi-classified organ donor Naive and $T_{CM}$ across all tissues.

Supplementary Table 15 **Differential expression of sorted $CD4^+$ and $CD8^+$ Naive vs $T_{CM}$** Differential expression of gene expression (GEX) between Naive and $T_{CM}$ sorted from PBMCs (Fig. 2), as identified by hashtag expression (see Methods).

Supplementary Table 16 **MMoCHi hierarchies (other multimodal datasets)** Subset definitions and thresholds used to construct MMoCHi hierarchies. Hierarchical relationships are defined by subset parents (subsets with no parent represent the first classification level). High-confidence events for each subset are selected as events matching the listed marker expression (above the positive threshold for all "Positive" markers, and below the negative threshold for all "Negative" markers). All gene expression markers are suffixed with "_gex". Thresholds for gene expression markers in log(CP100 + 1) for the Ab-seq and Xenium datasets and log(CP10k + 1) for the Glioma dataset. Thresholds for other modalities are in varied units, as described (see Methods).

Supplementary Table 17 **Important features for MMoCHi applied to other multimodal datasets** Gini impurity-based importance for each feature used for classification at each level of the MMoCHi hierarchies applied to the Ab-seq, Glioma, and Xenium datasets. Feature names for each level are suffixed with "_mod_GEX" for gene expression, "_mod_protein" for Ab-seq protein expression, "_mod_aneuploidy" for averaged chromosome expression in the Glioma dataset, and "_mod_physical_attributes" for morphological information extracted from Xenium imaging.
